# Single cell transcriptomics uncovers a non-autonomous *Tbx1*-dependent genetic program controlling cardiac neural crest cell deployment and progression

**DOI:** 10.1101/2022.08.01.502391

**Authors:** Christopher De Bono, Yang Liu, Alexander Ferrena, Aneesa Valentine, Deyou Zheng, Bernice E. Morrow

**Affiliations:** Department of Genetics, Albert Einstein College of Medicine, Bronx, NY, USA; Institute for Clinical and Translational Research, Albert Einstein College of Medicine, Bronx, NY, USA; Department of Neurology, Albert Einstein College of Medicine, Bronx, NY, USA; Department of Neuroscience, Albert Einstein College of Medicine, Bronx, NY, USA

**Keywords:** Cardiac neural crest cells, Outflow tract, Pharyngeal arch arteries, Tbx1, Cell fate progression, Cell-cell communication, Congenital heart disease

## Abstract

Disruption of cardiac neural crest cells (CNCCs) results in congenital heart disease, yet we do not understand the cell fate dynamics as these cells differentiate to vascular smooth muscle cells. Here we utilized single-cell RNA-sequencing of NCCs from the pharyngeal apparatus with heart in control mouse embryos and when *Tbx1*, the gene for 22q11.2 deletion syndrome, is inactivated. We uncovered three dynamic transitions of pharyngeal NCCs expressing *Tbx2* and *Tbx3* through differentiated CNCCs expressing cardiac transcription factors with smooth muscle genes, and that these transitions are altered non-autonomously by loss of *Tbx1*. Further, inactivation of *Tbx2* and *Tbx3* in early CNCCs resulted in aortic arch branching defects due to failed smooth muscle differentiation. Loss of *Tbx1* interrupted mesoderm to CNCC cell-cell communication with upregulation of BMP signaling with reduced MAPK signaling and failed dynamic transitions of CNCCs leading to disruption of aortic arch artery formation and cardiac outflow tract septation.

## Introduction

Neural crest cells (NCCs) are multipotent cells that migrate in three ordered streams from the rhombomeres in the neural tube to the pharyngeal apparatus where they differentiate to many cell types^1^. The pharyngeal apparatus is a dynamic embryonic structure consisting of individual pharyngeal arches (PA), forming in a rostral to caudal manner from mouse embryonic day (E) 8 to E10.5. A subset of pharyngeal NCCs migrate through the caudal pharyngeal arches, PA3-6, and surround the pharyngeal arch arteries (PAAs), while others continue to migrate to the cardiac outflow tract (OFT), both differentiating to vascular smooth muscle cells^2^. Ablation of NCCs from PA3-6 results in interruption of the aortic arch and arterial branching defects as well as persistent truncus arteriosus of the OFT^3^. These NCCs in PA3-6, are referred to as cardiac NCCs (CNCCs) based upon their position and known function in heart development as well as their differentiation to vascular smooth muscle. Understanding CNCC development is critical to determine the pathogenesis of human congenital heart defects such as those observed in 22q11.2 deletion syndrome (22q11.2DS) patients^4, 5^.

*TBX1,* encoding a T-box transcription factor, is the major gene for congenital heart disease in 22q11.2DS. Although 22q11.2DS is largely considered to be a neurocristopathy, *Tbx1* is not significantly expressed in CNCCs^6^, but it is strongly expressed in adjacent cells in the pharyngeal apparatus including the mesoderm. Global inactivation of *Tbx1* or conditional inactivation in the mesoderm using *Mesp1^Cre^* ^7^ in the mouse results in neonatal lethality with a persistent truncus arteriosus^8–10^, in part due to failed CNCC development^6^. Therefore, one of the main functions of *Tbx1* in the pharyngeal mesoderm is to signal to CNCCs to promote their development. In order to understand how CNCCs are affected non-autonomously in *Tbx1* mutant embryos, it is essential to define their transcriptional signatures and cardiac fate acquisition in the normal situation between E8.5 and E10.5, when *Tbx1* is expressed in the pharyngeal apparatus and when inactivated, on a single cell level.

Previously, single cell RNA-sequencing (scRNA-seq) of NCCs from early stages in the chick embryo identified expression of *Tgif1*, *Est1* and *Sox8* being important for early CNCC identity and fate decisions^11^. However, these were early migrating mesenchymal NCCs that also have the potential to contribute to the craniofacial skeleton and other cell types. Another seminal scRNA-seq study demonstrated that NCC fate choices are made by a series of sequential binary decisions in mouse embryos at E8.5-10.5^12^ but did not focus on detailed steps of cardiac fate acquisition or investigate *Tbx1* function^12^.

To uncover genetic signatures and dynamic transitions of CNCCs in the normal situation and when *Tbx1* is inactivated, we performed scRNA-seq of NCCs from control and *Tbx1* null mutant mouse embryos. We found that smooth muscle cell fate acquisition is in part dependent on two other T-box genes, *Tbx2* and *Tbx3*. When *Tbx1* is inactivated, we found failure of dynamic progression of CNCC maturation due to disruption of cell-cell communication from mesodermal cells, resulting in down regulation of MAPK signaling and upregulation of the BMP pathway, as well as affecting other, known and novel, ligand receptor interactions.

## Results

### Single cell transcriptional profiling of NCCs in the pharyngeal apparatus

We performed scRNA-seq of the *Wnt1-Cre, ROSA-EGFP* genetic lineage^13, 14^ in the mouse pharyngeal apparatus at E8.5, E9.5 and E10.5 (Fig. 1A-F). These stages correspond to developmental time points when *Tbx1* is highly expressed in cell types adjacent to NCCs. At E8.5, the anterior half of the embryo was dissected (Fig. 1A), while at E9.5 the pharyngeal apparatus with heart was microdissected (Fig. 1B). At E10.5 arches two to six and the heart were included in the dissection (Fig. 1C). EGFP positive NCCs were purified by FACS and the Chromium 10X platform was used to perform scRNA-seq and data from 36,721 NCCs were obtained (Supplementary Table 1). Unsupervised clustering was performed using Seurat software^15^ and individual clusters were identified (Fig. 1D-F).

**Figure 1:**
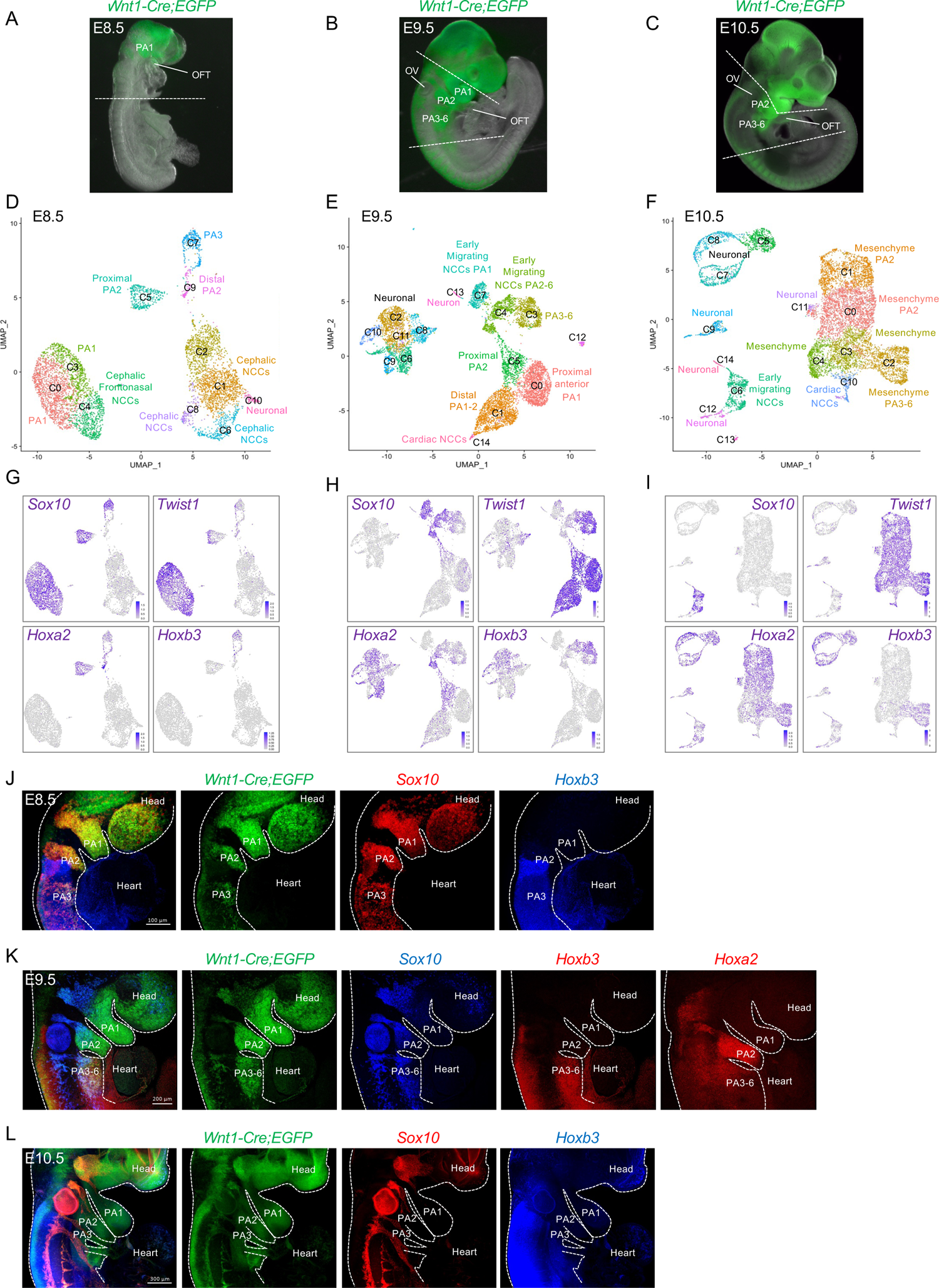
Single cell RNA-seq of NCCs from mouse embryos at E8.5-E10.5 reveals transcriptional heterogeneity within the pharyngeal region. A-C) *Wnt1-Cre;ROSA-EGFP* genetic lineage tracing shows the distribution of NCCs within the pharyngeal region and outflow tract of E8.5 (A), E9.5 (B) and E10.5 (C) embryos. The region rostral to the white dotted line of the embryo at E8.5 (A) and the pharyngeal region between the dotted lines in embryos at E9.5 and E10.5 including the heart were microdissected and EGFP positive NCCs were used for scRNA-seq. D-F) Seurat UMAP (Uniform Manifold Approximation and Projection) plots with cluster annotations of scRNA-seq data of NCCs at E8.5 (D), E9.5 (E) and E10.5 (F). (G-I) Expression of genes at E8.5 (G), E9.5 (H) and E10.5 (I) with highest expression in blue and lowest expression in gray. J-L) Wholemount RNAscope *in situ* hybridization of *Wnt1-Cre;ROSA-EGFP* embryos (n=3) at E8.5 (J), E9.5 (K) and E10.5 (L) with probes for *Sox10*, *Hoxb3* and *Egfp*, together and separated (colors are indicated above embryos). *Hoxa2* was examined at E9.5 as indicated. PA, pharyngeal arch; OFT, outflow tract; OV, otic vesicle. Scale bar: 100 μm in J, 200 μm in K and 300 μm in L.

Expression of *Sox10* and *Twist1* were used to identify early migratory and mesenchymal NCCs, respectively^16^. We used *Hox* and *Dlx* (Homeodomain) genes to provide spatial context to different arches (PA2-6; ^17^). At E8.5, *Sox10* and *Twist1* show overlap in expression, while at E9.5, expression became complementary, with a relative reduction of *Sox10*+ NCCs and increased *Twist1*+ NCCs in the expanded populations of mesenchymal NCCs (Fig. 1G-I). At E8.5 and E9.5, *Hoxa2* was expressed in PA2 and PA3-6, while *Hoxb3* was expressed only in PA3-6 containing NCCs that will invade the OFT and surround the PAAs (Fig. 1D,E,G,H). Using the *Hox* genes as a guide, at E9.5, early migrating *Sox10*+ NCCs of PA2 and PA3 were clustered together (cluster C4), suggesting that they have a similar transcriptional profile. At E10.5, the relative proportion of *Twist1* expressing mesenchymal cells increased with respect to reduction of *Sox10* expressing cells (Fig. 1F, 1I). Further, at E10.5, *Hoxa2* expression was expanded within the mesenchymal cell populations, and *Hoxb3* was expressed in NCCs of PA3-6 (cluster C3 at E9.5 is similar to C2 at E10.5; Fig. 1F, 1I). Additional marker genes are shown in Supplementary Data 1 (E8.5), 2 (E9.5) and 3 (E10.5). Spatial localization was confirmed for S*ox10*, *Hoxa2* and *Hoxb3* expression by wholemount RNAscope *in situ* hybridization (Fig. 1J-L). In addition, to anterior-posterior spatial localization of the cells, we identified their proximal-distal location in the PAs with *Dlx2*, *Dlx5* and *Dlx6* (Supplementary. Fig 1).

### Identification of cardiac NCC gene signatures

Differentiated NCCs of the OFT and PAAs express smooth muscle genes such as smooth muscle actin, *Acta2*^18, 19^. *Acta2* is a representative marker gene of smooth muscle cells that include expression of *Tagln*, *Myl9*, *Myh9*, and *Cnn1*. We identified a cluster of cells expressing *Acta2* at E9.5 and 10.5 (cluster C14, Fig. 2A; C10, Fig.3A), but not at E8.5. To delineate molecular signatures of cardiac NCCs (CNCCs), we evaluated genes that are co-expressed with *Acta2* and identified known genes for cardiac development including *Tbx2, Tbx3, Msx2, Isl1*, *Gata3* and *Hand2*, at E9.5 and E10.5 (Fig. 2A; Fig. 3A). These genes are not only expressed in *Acta2*+ cells but also in NCCs in the distal PA1-3 expressing *Dlx5* at E9.5 (C1, C3; Fig. 2A) and in PA2-6 at E10.5 (C2, C3, C4; Fig. 3A). At E9.5, we validated co-expression of ISL1 in CNCCs in the OFT of which some expressed ACTA2 (Fig. 2B). Further, RNAscope *in situ* analysis confirmed the expression of *Gata3*, *Isl1* and *Msx2* in CNCCs within the OFT at E9.5 (Fig. 2C-E). At E10.5, *Isl1* and *Gata3* were expressed in CNCCs within the cardiac cushions of the distal OFT and in the mesenchyme of the dorsal aortic sac wall and aortic sac protrusion (Fig. 3B). *Gata3* was expressed in a larger domain of the OFT than *Isl1*. When taken together, we now identify the genetic signatures of CNCCs of the forming OFT.

**Figure 2:**
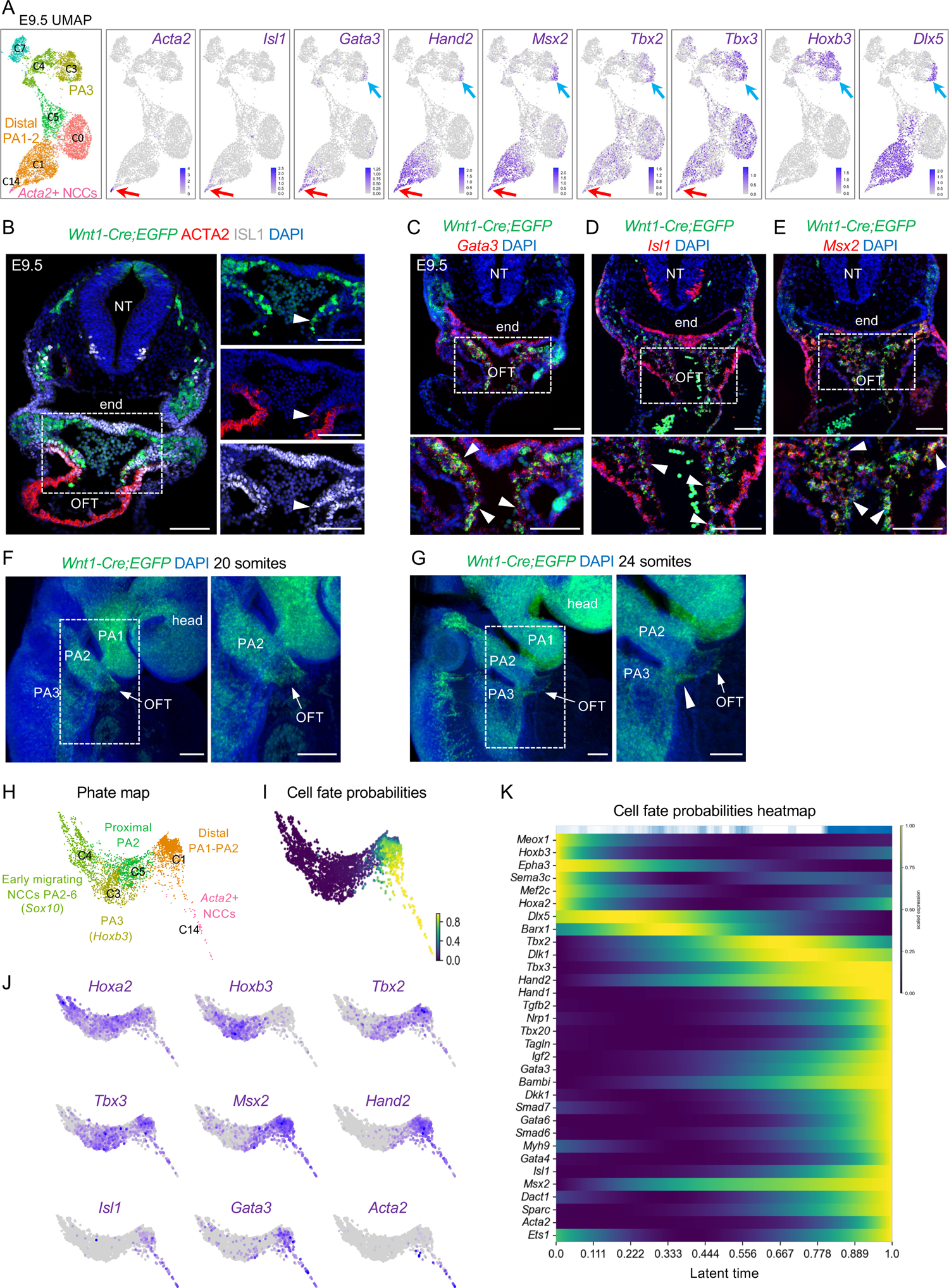
Transcriptional dynamics of cardiac NCCs at E9.5. A) UMAP plots of scRNA-seq data with genes that mark CNCCs identified by expression of *Acta2*, *Isl1*, *Gata3*, *Hand2*, *Msx2*, *Tbx2* and *Tbx3* with respect to *Hoxb3* (PA3) and *Dlx5* (distal PA1, 2, 3) expression. B) Immunostaining on traverse sections through *Wnt1-Cre;ROSA-EGFP* embryos showing Cre-activated EGFP (green), ACTA2 and ISL1 protein expression. Nuclei (blue) are labeled with DAPI. Arrowheads indicate CNCCs expressing ISL1 (n=3). C-E) RNAscope analysis of *Wnt1-Cre;ROSA-EGFP* embryos (n=3-4) for *Egfp*, *Gata3* (C), *Isl1* (D) and *Msx2* (E) expression. Nuclei (blue) are labeled with DAPI. Arrowheads indicate the expression of *Gata3*, *Isl1* and *Msx2* in CNCCS within the OFT. F,G) Wholemount RNAscope analysis of *Wnt1-Cre;ROSA-EGFP* embryos at 20 somites (F) and 24 somites (G) where the position of the OFT is indicated (white arrowhead). H) PHATE (Potential of Heat-diffusion for Affinity-based Transition Embedding) map of NCCs in clusters C1, C3, C4, C5 and C14 using Louvain clustering. I) PHATE map colored by cell fate probabilities, showing how each cell is likely to transition to CNCCs as defined by CellRank (yellow represent high cell fate probabilities). J) PHATE maps with expression of marker genes of CNCCs at different states of differentiation towards smooth muscle *Acta2* expressing cells. K) Heatmap from CellRank showing the expression of marker genes whose expression correlates with cardiac fate probabilities as latent time, with cells order by fate probabilities as latent time (see Supplementary data 4 for full list of genes). NT, neural tube; end, endoderm; OFT, outflow tract; PA, pharyngeal. Scale bars: 100μm.

**Figure 3:**
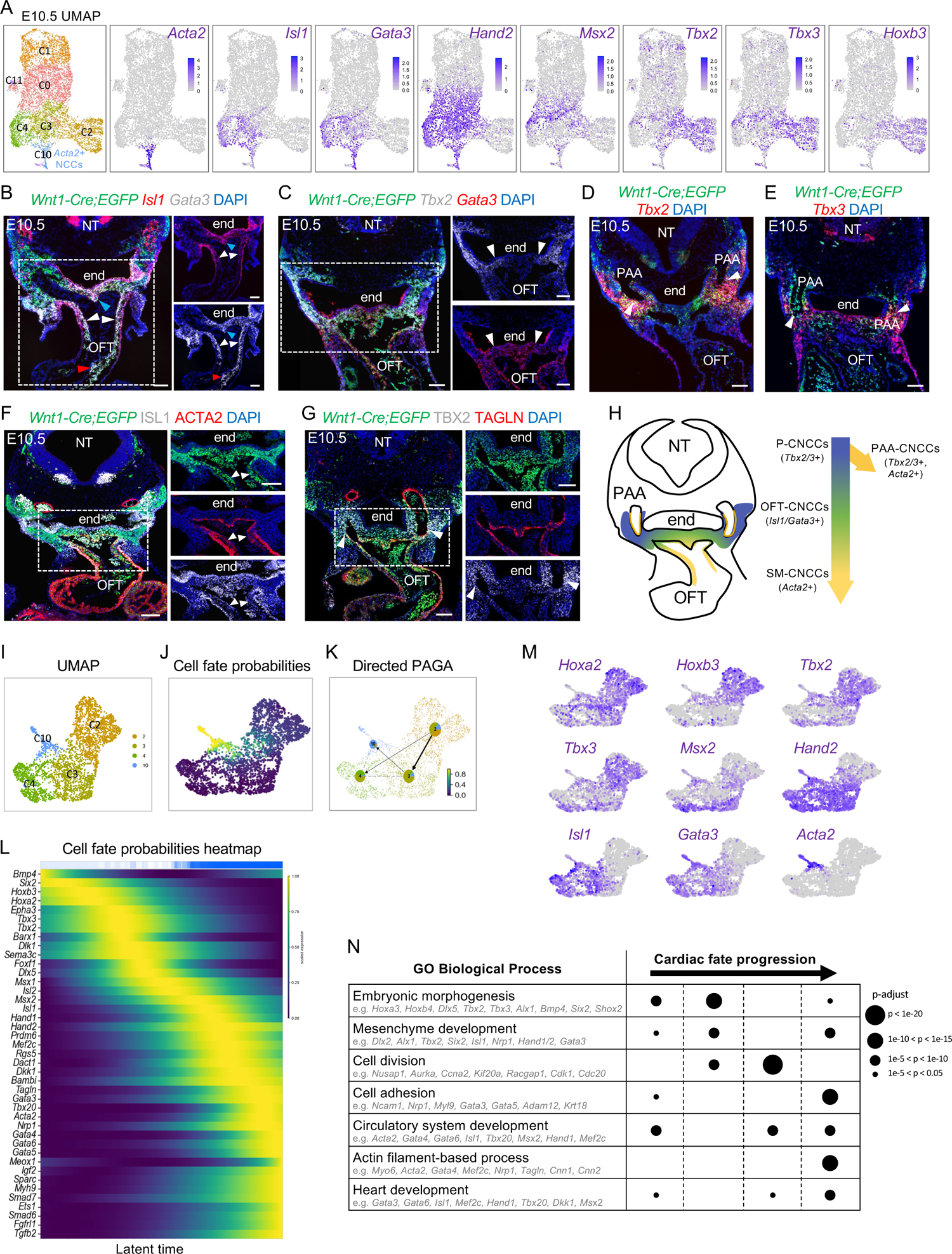
Transcriptional dynamics of cardiac NCCs at E10.5. A) Seurat UMP plots of scRNA-seq data from NCC populations with expression of marker genes. B-G) RNAscope *in situ* hybridization on traverse sections of *Wnt1-Cre;ROSA-EGFP* embryos (n=3-5) for *Egfp*, *Isl1* and *Gata*3 (B), *Egfp*, *Tbx2* and *Gata3* (C), *Egfp* and *Tbx2* (D) and *Egfp* and *Tbx3* (E) expression. Nuclei stained with DAPI are in blue. *Isl1* and *Gata3* are expressed in NCCs within of the OFT (white arrowheads in B) and at the level of the dorsal aortic sac wall and aortic sac protrusion (blue arrowheads in B). *Gata3* but not *Isl1* is expressed in the more proximal part of OFT (red arrowheads in B). *Tbx2* and *Gata3* expression overlaps in the dorsal wall of the OFT (white arrowheads in C). *Tbx2* and *Tbx3* are expressed in the mesenchyme dorsal to the aorta surrounding the PAAs at the level of the PA3-6 (white arrowheads in D and E). F, G) Immunostaining on traverse sections at the level of the OFT of *Wnt1-Cre;ROSA-EGFP* embryos (n=3) showing GFP, ISL1 and ACTA2 expression (F) and GFP, TBX2 and TAGLN expression (G). Expression of ISL1 is in smooth muscle cells of the OFT cushions (arrowheads in F) and expression of TBX2 is in smooth muscle cells of the PAAs (arrowheads in G). H) Schematic representation of a transverse section at the level of the OFT summarizing *Tbx2*, *Tbx3*, *Isl1*, *Gata3* and *Acta2* expression in CNCCs at E10.5. I) CellRank UMAP plot directed by RNA velocity and cell-cell similarity for clusters C2, C3, C4 and C10 from the feature plot in (A). J) UMAP plot for cell fate probabilities of CNCCs differentiating towards smooth muscle cells. K) Directed PAGA plot of NCCs. Pie charts show summarized fate probabilities of individual clusters, with blue representing the proportion of cells in each cluster with high probability to become smooth muscle expressing CNCCs of cluster, C10. L) Heatmap from CellRank showing the expression of selected genes whose expression correlates with transitions of CNCCs fate probabilities, with cells ordered by smooth muscle fate probabilities as latent time (see Supplementary data 5 for full list of genes). M) UMAP plots showing expression of marker genes in CNCCs. N) GO enrichment analysis of four groups of genes between *Bmp4* and *Gata6* (defined by their pseudotime; Supplementary data 6). Example of genes for each selected GO: biological processes are provided. Size of the dots indicate adjusted p-value (FDR B&Y). NT, neural tube; end, endoderm; PAA, pharyngeal arch artery. Scale bars: 100μm.

*Tbx2 and Tbx3* were widely expressed in pharyngeal NCCs at E9.5 (Fig. 2A), but their expression was restricted to cell clusters comprising PA3-6 at E10.5 (Fig. 3A). *Tbx2* and *Tbx3* were expressed immediately lateral and dorsal to *Isl1* and *Gata3* expressing CNCCs in embryos at E10.5 by RNAscope analysis (Fig. 3C-E). In addition, *Tbx2* and *Tbx3,* but not *Isl1* and *Gata3,* were expressed in NCCs surrounding the PAAs that are differentiating to smooth muscle at E10.5 (Fig. 3D,E,G). Both ISL1 and TBX2 proteins were expressed in smooth muscle cells of the OFT and PAAs, respectively (ACTA2 or TAGLN; Fig. 3F,G). Expression of *Tbx2*, *Tbx3*, *Isl1*, *Gata3* and *Acta2* in NCCs at E10.5 is illustrated in Fig. 3H. A subset of pharyngeal NCCs will form the CNCCs, defined as NCCs expressing markers specific to the cardiac or smooth muscle lineages. We refer to the CNCCs in the pharyngeal arches as P-CNCCs (Fig. 3H). Therefore, CNCCs can be subdivided into four populations based upon position and expression of cardiac or smooth muscle genes, referred to as P-CNCCs, PAA-CNCCs of PAAs expressing *Acta2*, OFT-CNCCs of the OFT expressing *Isl1* and *Gata3* and SM-CNCCs of the OFT that express *Acta2* (Fig. 3H).

We noted earlier that some CNCCs were located in clusters from PA1 and PA2 (C1; Fig. 2A), that are not typically considered to harbor CNCCs. Consistent with this, at E9.5 (20 somites) we found that the OFT was connected to PA2 and CNCCs from PA2 are entering the OFT (Fig. 2F). At late E9.5 (24 somites), the OFT was located between PA2 and PA3 and the first CNCCs from PA3 were entering the OFT (Fig. 2G). These data are consistent with evidence from a previous report^20^,that NCCs from anterior arches also contribute to the developing heart.

### Cardiac NCC fate dynamics that drive differentiation to smooth muscle cells

To uncover CNCC cell fate dynamics at E9.5, we used CellRank software^21^ (Fig. 2H, I). We discovered genes that were progressively activated during the transition from pharyngeal NCCs to SM-CNCCs, which are candidate cardiac lineage driver genes (Fig. 2J, K; Supplementary data 4 for the full list of genes). Our analysis indicates that CNCCs progressively activate *Tbx2*, *Tbx3*, *Msx2, Hand2*, *Gata3 and Isl1* expression during their commitment towards *Acta2+* smooth muscle cells at E9.5 (Fig. 2J,K).

To understand how CNCCs progress at E10.5, when there are more smooth muscle cells in the pharyngeal arches, we used CellRank software and generated PAGA (partition-based graph abstract) plots (Fig. 3I-K). The cell fate probability map from CellRank identified cells with a high potential to differentiate to smooth muscle fates (from cluster C2 and C3 to C10; Fig. 3J). The PAGA plots further indicated that some pharyngeal NCCs (cluster C2) are P-CNCCs and they transition to OFT-CNCCs (cluster C3) that then transition to SM-CNCCs (cluster C10; blue color fraction in pie chart; Fig. 3K). This data also indicates that a small fraction of P-CNCCs may directly differentiate to smooth muscle cells (blue fraction in the pie chart in C2), in agreement with *Tbx2* and *Tbx3* expression in PAA-CNCCs at E10.5 (Fig. 3D, E, G). We identified genes whose expression correlates with SM-CNCC fate acquisition (Fig. 3L, M; Supplementary data 5 for full list of genes at E10.5). Representative genes were ordered according to their expression peak in pseudotime and included *Tbx2*, *Tbx3*, *Foxf1*, *Isl2*, *Msx2*, *Isl1*, *Hand1*, *Hand2*, *Mef2c*, *Rgs5*, *Gata3*, *Acta2*, *Gata4* and *Gata6*.

We generated lineage driver gene sets by dividing the genes in the fate probabilities heat map from *Bmp4* to *Gata6*, least to most differentiated to SM-CNCCs, to four groups of equal size (Fig. 3L; Supplementary data 5). Next, we performed Gene Ontology (GO) enrichment analysis using ToppGene Suite^22^ to understand the function of the genes in each group (Fig. 3N; Supplementary data 6). Our analysis indicates that the initially activated genes of pharyngeal NCCs that include some P-CNCCs, are associated with general pharyngeal arch development processes (e.g. *Hox*, *Dlx*, *Six2* genes). Then cell division (cell cycle) genes are highly expressed, consistent with the expansion of pharyngeal NCCs during development^23^, together with cardiac development genes. Finally, genes important in cardiac development, cell adhesion and actin-filament processes (*Hand1, Gata3/4/5/6, Isl1, Acta2*) become strongly expressed (Fig. 3N). In addition, our functional enrichment analysis identified genes associated with congenital heart disease such as tetralogy of Fallot, double outlet right ventricle and ventricular septal defects (Supplementary data 6), which supports the importance of the genetic program of CNCCs in OFT formation and disease.

Thus, here we identified a specific CNCC transcriptomic signature at E9.5-10.5 and revealed that cell fate acquisition to smooth muscle cells requires a multistep specification process. We additionally identified new genes such as *Dkk1*, *Gata3*, *Foxf1*, *Isl2*, *Tbx2*, *Tbx3*, *Rgs5,* among others, which have not yet been considered as CNCC markers (Supplementary data 4 and 5).

### *Tbx2* and *Tbx3* are required in cardiac NCCs for aortic arch branching

*Tbx2* and *Tbx3* are expressed in multiple tissue types within the pharyngeal apparatus and global inactivation of both genes leads to early embryonic lethality with severe cardiac defects^24–27^. To understand the requirement of *Tbx2* and *Tbx3* in NCCs, we generated *Wnt1-Cre/+;Tbx2^f/f^;Tbx3^f/f^*double conditional mutant embryos (*Tbx2/3* cKO). We performed intracardiac ink injection and histological analysis at E15.5 (Fig. 4A-I) and found that 38.5% of *Tbx2/3* cKO embryos had an aberrant retro-esophageal right subclavian artery (ARSA) but no intracardiac defects (Fig 4A-I and M). No defects were identified in *Wnt1-Cre/+;Tbx2^f/f^;Tbx3^f/+^* nor in *Wnt1-Cre/+;Tbx2^f/+^;Tbx3^f/f^* embryos. The right subclavian artery is formed from the right 4^th^ PAA. By immunostaining on coronal sections of *Wnt1-Cre/+;Tbx2^f/f^;Tbx3^f/f^;ROSA-EGFP^f/+^*embryos at E11.5 using GFP and ACTA2 antibodies, we found that NCCs contributed to the right 4^th^ PAAs but failed to differentiate into smooth muscle cells (Fig. 4J-L). *Bmp4* and *Foxf1* have been identified as regulators of smooth muscle cell differentiation in other organs^28^. We found that *Bmp4* and *Foxf1* expression is activated temporally after *Tbx2* and *Tbx3* expression during cardiac fate acquisition (Fig. 3L).

**Figure 4:**
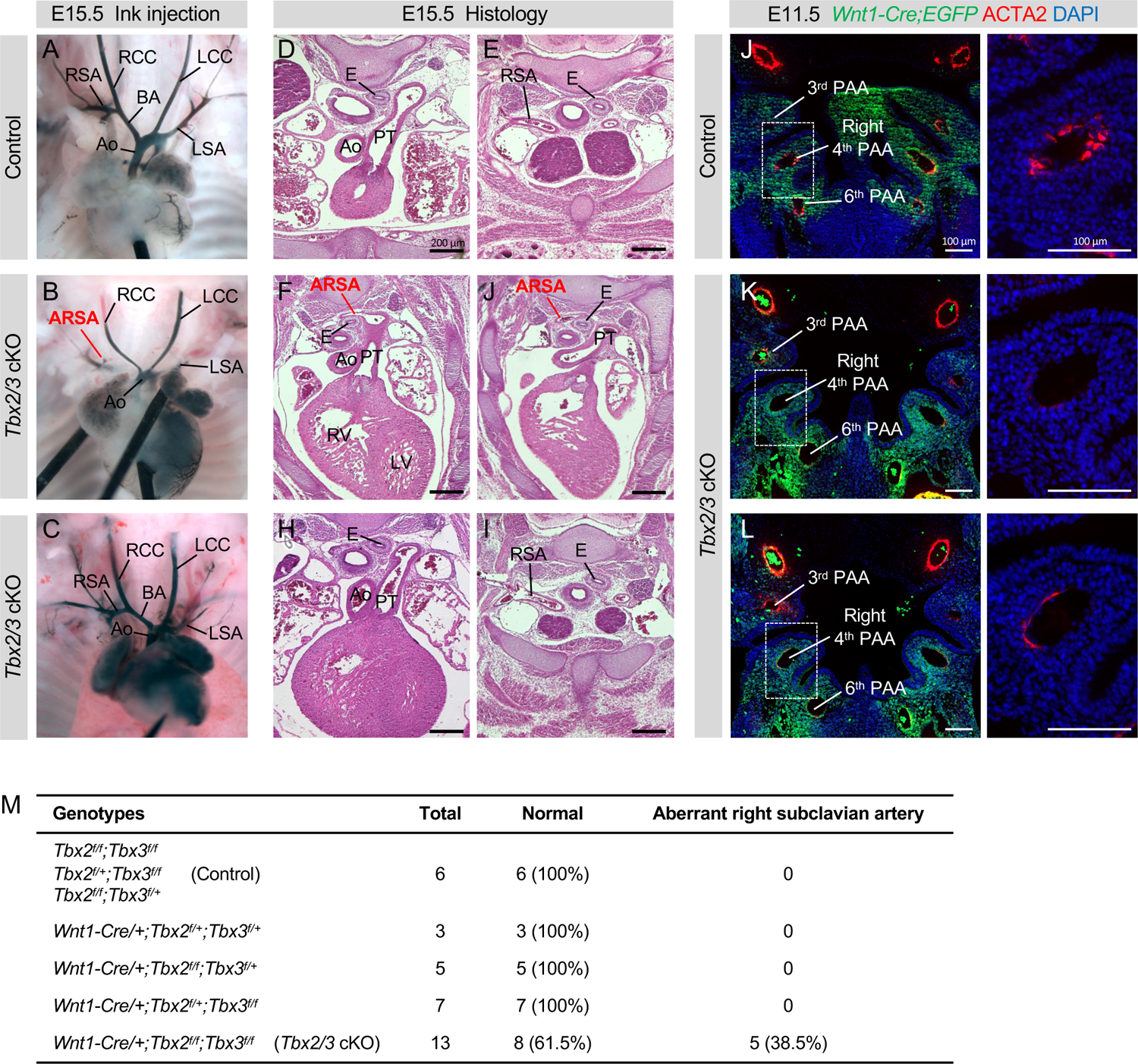
*Tbx2* and *Tbx3* are required together for arterial branching from the aortic arch. A-D) Intracardiac ink injection of control (A) and *Wnt1-Cre;Tbx2^f/f^;Tbx3^f/f^* conditional null mutant embryos at E15.5 (B,C). Double *Tbx2/Tbx3* cKO embryos have a partially penetrant aberrant retro-esophageal right subclavian artery (ARSA) (B). D-I) Haematotoxin and Eosin staining on traverse sections of control (D,E) and *Tbx2/Tbx3* cKO mutant embryos (F-I) at E15.5. Panels F and J show a *Tbx2/3* cKO embryo with ARSA and panels H and I shows a *Tbx2/3* cKO embryo with a normal right subclavian artery. J-K) Immunofluorescence on coronal sections of controls at E11.5 (I) and *Tbx2/3* cKO embryos (K,L), at the level of the pharyngeal arch arteries for GFP and ACTA2 expression. Nuclei are stained with DAPI. Right panels are high magnification of the dashed regions in J, K and L. Note the strong reduction of ACTA2 expression in the right 4^th^ PAA in *Tbx2/3* conditional mutant embryos compared to control embryos. M) Table of the cardiovascular defects in *Tbx2/Tbx3* conditional mutants. RSA, right subclavian artery; RCC, right common carotid; LCC, Left common carotid; BA, brachiocephalic artery; Ao, aorta; LSA, left subclavian artery; PT, pulmonary trunk; E, esophagus; PAA, pharyngeal arch artery.

### Disruption of cardiac NCCs by loss of *Tbx1*

In *Tbx1* null mutant embryos, the caudal pharyngeal apparatus is hypoplastic and unsegmented at E9.5 and E10.5 due in part to failed deployment of NCCs^6^ (Fig. 5A, Fig 7A). Further, CNCCs fail to enter the shortened cardiac OFT, leading to a persistent truncus arteriosus later in development^6^. We found that *Tbx1* was not noticeably expressed in NCCs (Supplementary Fig. 2) and its conditional deletion in NCCs using *Wnt1-Cre* did not lead to cardiac defects (Supplementary Fig. 3).

**Figure 5:**
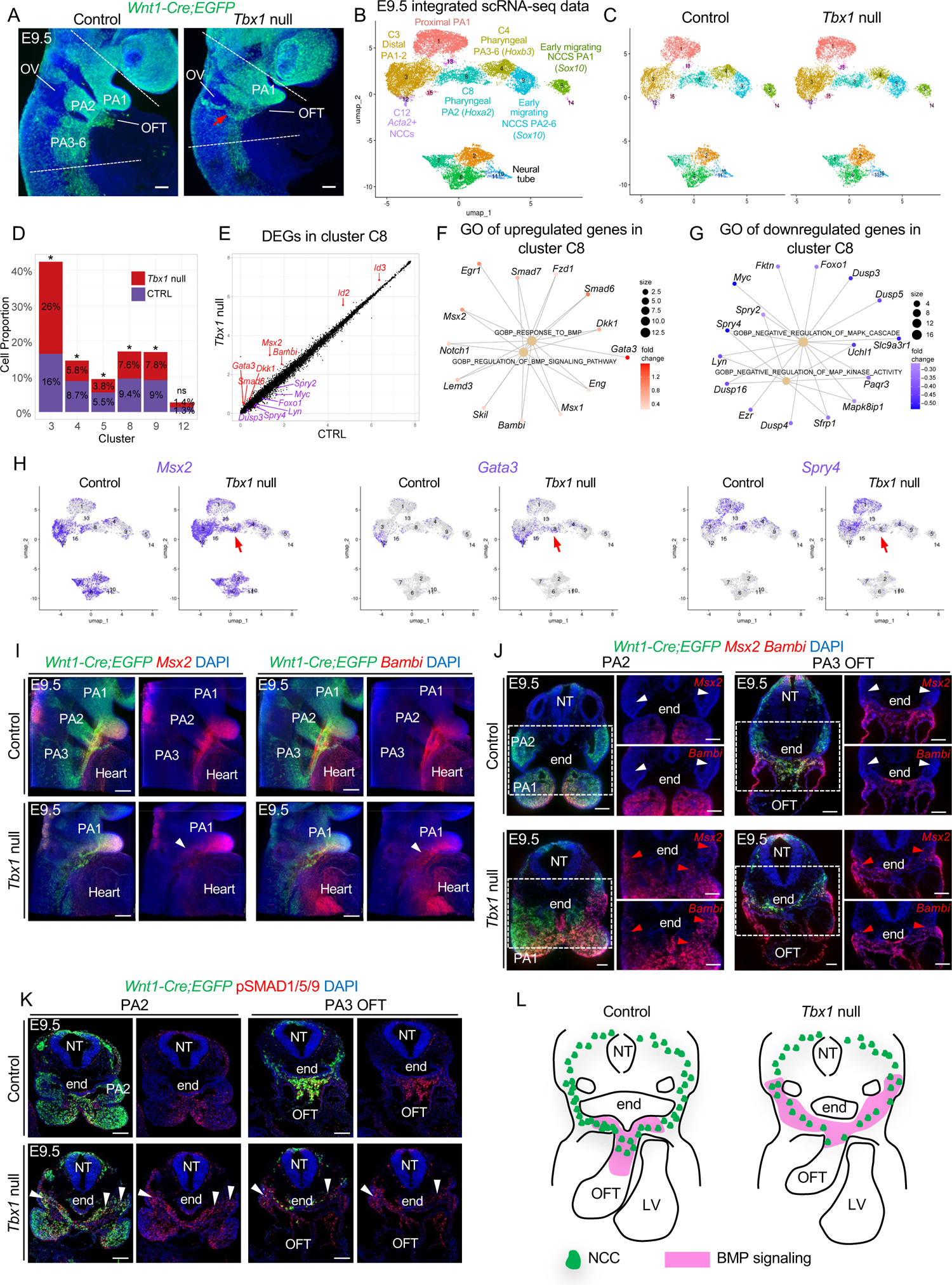
scRNA-seq identifies upregulation of the BMP pathway by inactivation of *Tbx1* at E9.5. A) *Wnt1-Cre;ROSA-EGFP* lineage tracing (green) shows mis-localization and reduced number of NCCs within the pharyngeal region of *Tbx1* null embryos (red arrow). DAPI is in blue. NCCs from the region between the two dashed lines were used for scRNA-seq. B) RISC UMAP plot of integrated scRNA-seq data from NCCs of control and *Tbx1* null embryos. C) UMAP plots colored by clusters from control (left) and *Tbx1* null embryos (right). D) Stack bar graph shows proportion of NCCs in indicated clusters, computed as the number of cells in these clusters divided by total number of cells in control or *Tbx1* null embryos. Two proportion z test was used to evaluate cell proportion differences between control and *Tbx1* null embryos (cluster C3: P-value = 1.14e-56; C4: P-value = 9.24e-15; C5: P-value = 1.46e-8; C8: P-value = 1.23e-5; C9: P-value = 4.32e-3; C12: P-value: 0.66) (*, P-value < 0.05%; ns, not significantly different) E) Scatter plot shows differential gene expression in cluster C8 from control and *Tbx1* null embryos at E9.5. Representative DEGs in BMP (red) and MAPK (purple) pathways are indicated. F-G) GO enrichment analysis using clusterProfiler for upregulated (F) and downregulated (G) genes in C8 of *Tbx1* null embryos. Note upregulation of genes involved in the BMP pathway and downregulation of genes in the MAPK pathway. The Category Netplots links genes with Gene Ontology terms. H) UMAP plots show expression level (purple) of *Msx2*, *Gata3* and *Spry4* in NCCs split by control and *Tbx1* null embryos. I) *Msx2*, *Bambi* and *Egfp* wholemount RNAscope *in situ* hybridization with DAPI of control and *Tbx1* null embryos (n=3-7). The arrowheads indicate ectopic expression of *Msx2* and *Bambi* in *Tbx1* null embryos. J) *Egfp*, *Msx2* and *Bambi* RNAscope assays on transverse sections of control and *Tbx1* null embryos (n=3) showing ectopic and dorsal expression of *Msx2* and *Bambi* in migrating NCCs (red arrowheads). *Msx2* and *Bambi* expression are absent in the proximal part of the PA2 and PA3 of control embryos (white arrowheads). K) Fluorescent immunostaining for GFP and P-SMAD1/5/9 on transverse sections of control (n=5) and *Tbx1* null (n=4) embryos shows increase in P-SMAD1/5/9 in NCCs of PA2-3 (arrowheads). L) Schematic representation of control (left) and *Tbx1* null (right) sections at E9.5 showing reduced CNCCs contributing the shortened OFT and ectopic posterior BMP signaling. PA, pharyngeal arch; OV, otic vesicle; OFT, outflow tract; NT, neural tube; end, endoderm. Scale bars: 100μm.

To understand how the absence of *Tbx1* affects development of CNCCs, we performed scRNA-seq of NCCs isolated from the microdissected pharyngeal region plus heart of *Tbx1* null mutant embryos at E9.5. We obtained sequencing data from 11,301 NCCs (Fig 5A; Supplementary Table 1) and integrated scRNA-seq data from control and *Tbx1* null embryos using RISC (Robust Integration of scRNA-seq) software^29^. Even though there were visibly fewer NCCs in the pharyngeal apparatus (Fig. 5A), there were no missing cell clusters in *Tbx1* null embryos (Fig. 5B-C). We then compared the proportion of cells in each cluster among the total number of NCCs in each dataset. As expected, there was a reduction in the relative proportion of NCCs from *Tbx1* null embryos as compared to controls, as shown in Fig. 5D, in PA3 (C4,1.5 fold), in proximal PA2 (C8, 1.4 fold), in cluster C9 corresponding early migrating NCCs in PA2 and PA3 (1.2 fold), and in C5 corresponding to early migrating NCCs in PA1 (1.4 fold). In addition, there was an increase of the relative proportion of NCCs in cluster C3 (1.6 fold) that corresponds to the distal part of PA1 and PA2 (Fig. 5D) and it is known that cells from PA2 abnormally migrate to PA1 in *Tbx1* null embryos^6^.

### Altered BMP and MAPK signaling pathways in the absence of *Tbx1*

We examined the data to identify differentially expressed genes (DEGs) in mutant versus control embryos at E9.5. Surprisingly, we found few DEGs per cluster at this stage (Supplementary data 7). However, in *Tbx1* null embryos there was a clear increase in the expression of genes that act downstream of BMP signaling in proximal PA2 and PA3 (clusters C4, C8; Fig. 5E). This includes increased expression of *Msx2*, *Bambi*, *Gata3*, *Dkk1*, *Smad6*, *Id2* and *Id3*. Our analysis also revealed a downregulation of the expression of genes in the MAP kinase (mitogen-activated protein kinase) signaling pathway including *Spry2*, *Spry4*, *Myc*, *Foxo1*, *Lyn*, and *Dusp3* (Fig 5E). Signaling by BMP^30^ and growth factors activating the MAPK pathway^31^ are two signaling pathways known to be critical for NCC development and migration during embryogenesis. Gene enrichment analysis of DEGs by cluster profiler R software^32^ confirmed an increase in expression of genes in the BMP signaling pathway (Fig. 5F, Supplementary data 8) and a decrease in expression of genes related to negative regulation of MAPK cascade and activity (Fig. 5G, Supplementary data 9). There was an increase in expression of *Msx2* and *Gata3* (downstream in the BMP pathway) as well as reduced expression of *Spry4* (MAPK pathway) in scRNA-seq data of *Tbx1* null embryos (Fig. 5H). Expression of BMP downstream genes, *Msx2* and *Bambi*, were expanded dorsally in *Tbx1* null mutant embryos by wholemount RNAscope *in situ* and 3D reconstruction (Fig. 5I). These results were confirmed by RNAscope assays on traverse sections of control and *Tbx1* null embryos (Fig. 5J). In addition, there was an increase and ectopic expression of P-SMAD1/5/9, marking an increase in BMP signaling, in NCCs towards the dorsal part of the pharyngeal apparatus in *Tbx1* null embryos (PA2, PA3; Fig. 5K). These data suggest that altered BMP and MAPK signaling might affect NCC development in *Tbx1* null embryos. A schematic representation of expanded BMP signaling and reduced NCCs migrating to the shortened OFT in *Tbx1* null embryos at E9.5 is shown in Fig. 5L.

### Cell-cell communication from the mesoderm to NCCs is disrupted in the absence of *Tbx1*

During formation of the heart, NCCs receive critical signaling from adjacent mesodermal cells (Fig. 6A). We investigated cell-cell communication and how it is disrupted in the absence of *Tbx1* at single cell resolution using CellChat software^33^.

**Figure 6:**
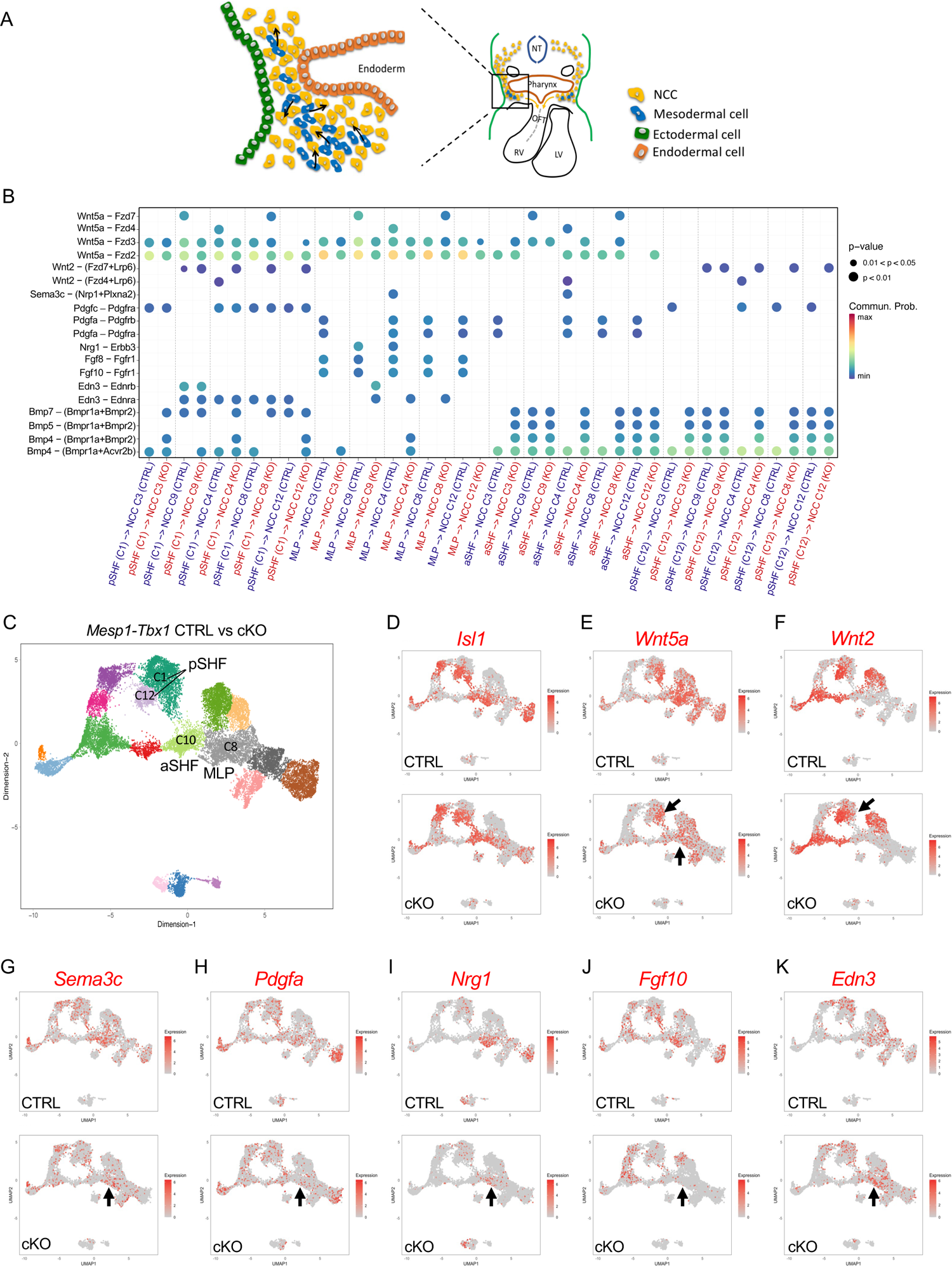
Cell-cell communication from the mesoderm to NCCs is altered in the absence of *Tbx1* at E9.5. A) Schematic representation of a transverse section showing signaling (arrows) from pharyngeal mesoderm cells (blue) to NCCs (yellow) in the caudal pharyngeal apparatus. B) Bubble plots show representative cell-cell signaling from *Mesp1^Cre^* derived mesodermal cells to NCCs that were significantly altered (p-value indicated by size of dot and color; right) in *Tbx1* mutant embryos. Each dot represents a ligand-receptor pair interaction (Y-axis) between a specific cluster in the mesoderm cells and NCCs (X-axis). Clusters include anterior and posterior second heart field (aSHF, pSHF) and the multilineage progenitors (MLP). C) UMAP from integration of two replicates of *Mesp1^Cre^;ROSA-EGFP* (CTRL) and *Mesp1^Cre^;Tbx1^f/f^;EGFP* (*Tbx1* cKO) datasets. D) UMAP plots showing *Isl1* expression in control and *Tbx1* cKO cells. E-K) UMAP plots showing the expression of ligand genes including *Wnt5a* (E), *Wnt2* (F), *Sema3c* (G), *Pdgfa* (H), *Nrg1* (I), *Fgf10* (J) and *Edn3* (K) in *control* and *Tbx1* cKO embryos. Arrows indicate cell clusters with gene expression changes in *Tbx1* cKO embryos.

Inactivation of *Tbx1* in the mesoderm results in similar pharyngeal hypoplasia and altered NCC distribution as in global null embryos, implicating the pharyngeal mesoderm as being critical to signal to NCCs^34^. To identify *Tbx1*-dependent signals from the mesoderm to NCCs, we investigated existing scRNA-seq data from *Mesp1^Cre^* control and *Tbx1* conditional null embryos at E9.5^35^. We focused on mesodermal subpopulations, expressing *Tbx1*, that are adjacent to the NCCs including the anterior and posterior second heart field (aSHF; pSHF)^36, 37^. We also included a critical *Tbx1*-dependent multilineage progenitor population (MLP) in the pharyngeal mesoderm required for cell fate progression to the aSHF and pSHF^35^. We examined signaling to NCCs in clusters corresponding to migrating NCCs of the future PA2-6 (C9), distal part of PA1-2 (C3), mesenchyme of PA2 (C8) and PA3-6 (C4), and CNCCs of the OFT (C12) in integrated scRNA-seq data from control and *Tbx1* null embryos at E9.5 (Fig. 5; Fig. 6B). Representative results of ligand-receptor pairs altered when *Tbx1* is inactivated are shown in Fig. 6B, and the complete set of pairs are in Supplementary Fig. 4.

Affected ligands in the mesoderm include *Wnt5a, Wnt2, Sema3c, Pdgfa Nrg1, Fgf8, Fgf10, Bmp4* and *Edn3* and others. To validate relationships, we analyzed integrated *Mesp1^Cre^* data (Fig. 6C; MLPs-C8, aSHF-C10 and pSHF-C1+C12). *Isl1*, is a critical gene required for OFT development^38^, and it is expressed in the MLPs, aSHF and pSHF (Fig. 6D). We examined expression changes of *Wnt5a*, *Wnt2*, *Sema3c*, *Pdgfa*, *Nrg1*, *Fgf10* and *Edn3* (Fig. 6E-K). These genes were altered in expression in the cell types specified and, in the direction of altered signaling (decreased or increased in the mutant embryos), as indicated in Fig. 6B.

Reduced expression of *Wnt5a*^39, 40^*, Fgf8*^35, 41, 42^*, Fgf10*^43, 44^, *Sema3c*^45^ and *Nrg1*^35^ ligands and increase of *Wnt2*^46^ are consistent with previous *in vivo* studies of *Tbx1* mutant embryos, however these were not known with respect to cell-cell communication to NCCs. With this data, we show on a single cell level, that this signaling to NCCs is altered in *Tbx1* mutant embryos. Two additional ligand genes, *Edn3* and *Pdgfa* were not investigated regarding *Tbx1* (Fig. 6H, 6K). *Edn3* encodes an endothelin ligand important in cell migration but not well known with respect to *Tbx1*. *Pdgfa* encodes a growth factor regulating cell survival, proliferation and migration and PDGF signaling is required in NCC development^47^.

We investigated CellChat results for the BMP and MAPK pathways that were altered in NCCs when *Tbx1* was inactivated. An abnormal increase of the BMP signaling from the mesoderm to NCCs in *Tbx1* mutant embryos was found through *Bmp4/5/7* ligands (Fig. 6B). *Fgf8* and *Fgf10* are ligands in FGF signaling that act through the MAPK pathway and it is well known that they are reduced in expression in *Tbx1* mutant embryos^35, 41–44^ (Fig. 6J). Similarly, we found that *Fgf8* and *Fgf10* were reduced in expression in the mesoderm and signaling to NCCs was altered (Fig. 6B and Supplementary Fig. 4). It is known that FGF and BMP pathways can act antagonistically^48, 49^. Therefore, it is possible that reduction of FGF and changes in other ligands in the adjacent mesoderm, could result in ectopic BMP and reduced MAPK signaling in NCCs leading to their failed progression.

### Failed cardiac cell fate progression of NCCs in the absence of *Tbx1* at E10.5

To further understand how contribution of NCCs to the OFT is altered in the absence of *Tbx1*, we performed scRNA-seq of NCCs in *Tbx1* null embryos at E10.5, when the caudal pharyngeal apparatus is extremely hypoplastic (Fig. 7A). We integrated scRNA-seq data from two control replicates (21,561 cells) and two *Tbx1* null replicates (17,840 cells) using RISC software (Fig. 7B). The integrated datasets clearly show a strong reduction in the number of NCCs in most clusters in *Tbx1* null embryos including OFT-CNCCs (*Isl1/Gata3*+; C10) and SM-CNCCs (*Acta2+*; C13) as shown in Fig. 7C, except that the relative proportion of pharyngeal NCCs that contain P-CNCCs (*Tbx2/Tbx3+*; C4) is not changed, after considering the total cells being sequenced (Fig. 7D). Our analysis showed that the cell fate probabilities point from pharyngeal NCCs containing P-CNCCs (C4) and OFT-CNCCs (C10) toward SM-CNCCs (C13) as shown in Fig. 7E. We generated a list of DEGs in clusters C4 and C10, between control and mutant embryos at E10.5 (Supplementary data 10). Consistent with data from E9.5, *Msx2*, *Bambi*, *Gata3* and *Dkk1* were increased and ectopically expressed in pharyngeal NCCs in *Tbx1* null embryos (Fig. 7F, Supplementary data 10), suggesting an abnormal upregulation of BMP signaling in the absence of *Tbx1*, consist with data in Figure 6. By GO analysis, we found downregulation in expression of genes involved in embryonic organ development and mesenchyme development in pharyngeal NCCs that contain P-CNCCs (Fig. 7G; Supplementary data 11), suggesting dysregulation of NCC development. Interestingly, there was an upregulation of genes that inhibit cell cycle progression of OFT-CNCCs (Fig. 7H; Supplementary data 12). By immunostaining, we confirmed the overall reduction in the number of CNCCs within the OFT and NCCs in the pharyngeal region of *Tbx1* null embryos (Fig. 7I,J). We also confirmed a reduced number of ISL1+ NCCs in dorsal aortic sac wall mesenchyme and distal OFT and absence of the aortic sac protrusion in *Tbx1* null embryos (Fig. 7I). Supporting the scRNA-seq data, immunostaining experiments indicated that TBX2 (and likely TBX3) expression is maintained in NCCs located in the lateral part of the pharyngeal apparatus. We also noticed normal differentiation of the CNCCs within the OFT of *Tbx1* null embryos despite that there are fewer cells (Fig. 7J). Together this suggests a failure of cardiac fate progression between pharyngeal NCCs and OFT-CNCCs states in the absence of *Tbx1*.

**Figure 7:**
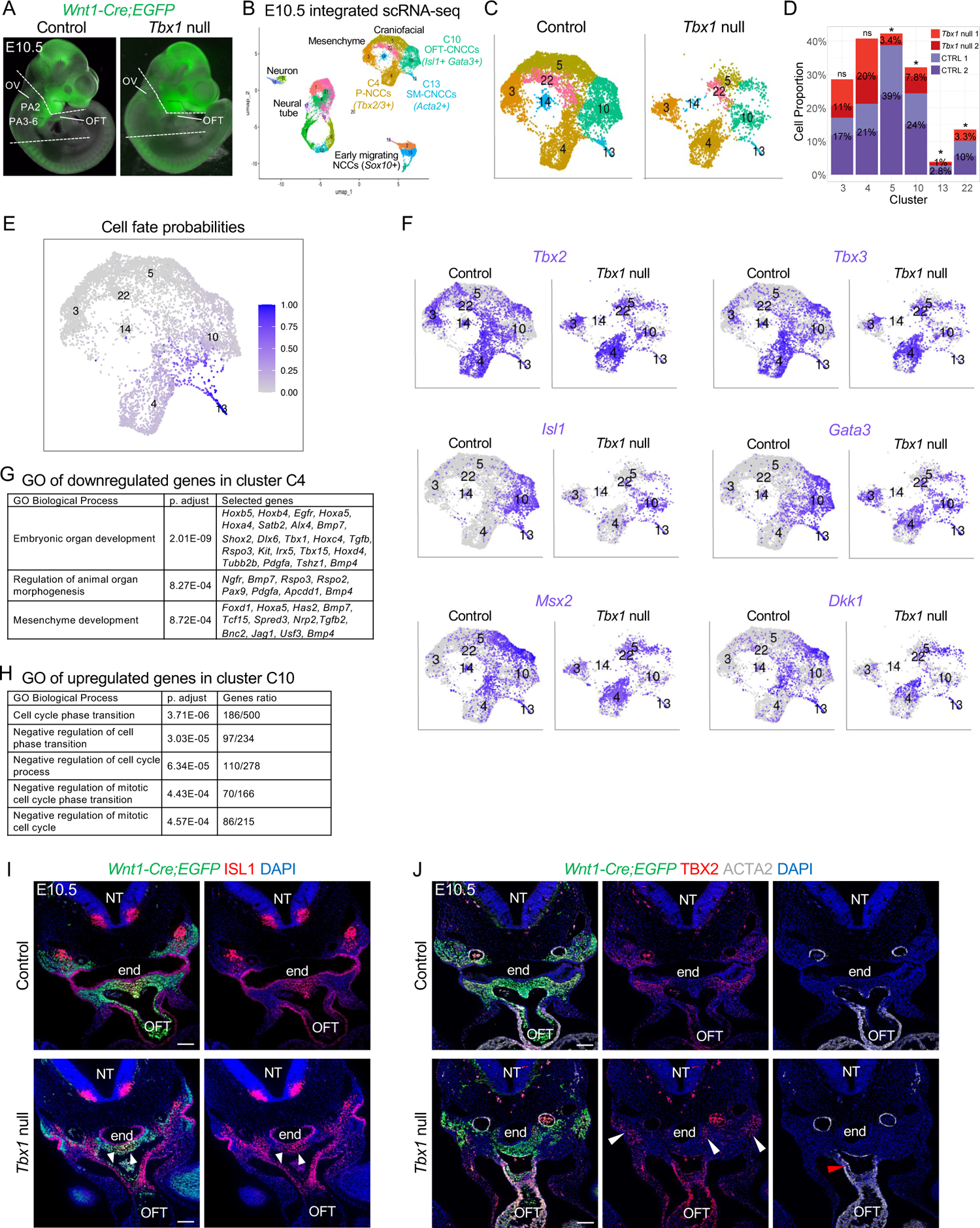
Failure of CNCC cell fate progression by loss of *Tbx1* at E10.5. A) The NCCs between the dotted lines and heart in control and *Tbx1* null embryos were used for scRNA-seq. B) RISC UMAP plot of integrated scRNA-seq data from NCCs of two replicates of control and *Tbx1* null embryos. C) UMAP plots of pharyngeal and heart clusters from control (left) and *Tbx1* null (right) embryos. D) Stack bar graph shows proportions of NCCs in selected clusters divided by the total cell number of cells in control or *Tbx1* null replicates. The proportion of NCCs from *Tbx1* null embryos in cluster C10 is strongly reduced but not in C4. Two proportion z test was used to evaluate cell proportion differences between control and *Tbx1* null embryos. (*, P-value < 0.05%; ns, not significantly different). E) UMAP plot colored by cell fate probabilities. F) UMAP plots showing expression level of *Tbx2*, *Tbx3*, *Isl1*, *Msx2*, *Gata3*, *Dkk1* in NCCs from control or *Tbx1* null embryos in pharyngeal NCC clusters. *Msx2*, *Gata3* and *Dkk1* show increased expression in cluster C4. G) GO analysis for downregulated genes in C4. H) GO analysis of upregulated genes in C10. I) Fluorescent immunostaining for GFP (green) and ISL1 (red) with DAPI (blue) on transverse sections of control (n=4) and *Tbx1* null (n=3) embryos. There is reduced ISL1 positive NCCs (arrowheads). J) Fluorescent immunostaining for GFP (green), TBX2 (red), and ACTA2 (grey) on transverse sections of control (n=4) and *Tbx1* null (n=3) embryos, DAPI is in blue. Expression of TBX2 in NCCs is lateral to the pharyngeal endoderm (end; white arrowheads). There are fewer CNCCs expressing ACTA2 within the OFT of *Tbx1* null embryos (red arrowhead). NT, neural tube; end, endoderm; OFT, outflow tract. Scale bars: 100μm

## Discussion

In this report, we identified the signatures and cell fate dynamics of CNCCs. We focused on pharyngeal NCCs in the mouse at developmental stages when *Tbx1*, the gene for 22q11.2DS is highly expressed and functions. We determined the mechanisms by which *Tbx1* non-autonomously regulates CNCC maturation at a single cell level and found altered BMP and MAPK signaling may contribute to cardiovascular malformations when *Tbx1* is inactivated. We also uncovered novel genes and ligand-receptor pairs with respect to cell-cell communication from mesoderm to NCCs that are new to our understanding of CNCC fate progression.

NCCs are multipotent and differentiate to many cell types including smooth muscle cells. Through examination of transitional dynamics along with embryonic localization by *in situ* analysis, we uncovered three main transition states from pharyngeal to smooth muscle expressing CNCC derivatives, termed P-CNCCs, OFT-CNCCs and SM-CNCCs as shown in the model in Fig. 8A. The P-CNCCs express *Hox* and *Dlx* genes, as well as those implicated in cardiac development including *Tbx2*, *Tbx3*, *Six2*, *Shox2*, *Bmp4*, *Prdm1*, *Daam2*, *Scube1*, *Angptl1*, *Tfap2b* and *Mef2c.* Our data also suggests that some *Tbx2/3*+ cells differentiate directly into *Acta2*+ smooth muscle in the PAAs (PAA-CNCCs; Fig. 8A). A role of *Tbx2* and *Tbx3* in CNCCs in PAA development have not been previously described and we found that inactivation results in abnormal arterial branching at reduced penetrance, indicating besides function as markers of P-CNCCs, they have a role in smooth muscle differentiation. The second state of OFT-CNCCs express cardiac transcription factors such as *Hand1/2, Msx1/2, Mef2c* and *Gata3* as well as *Isl1* and *Isl2* (Fig. 8A). These cells are required for OFT development, as determined by conditional inactivation studies in NCCs of *Hand2*^50^, *Msx1* and *Msx2*^51^. Expression of *Isl1* in CNCCs contributing to the OFT is consistent with a dual lineage tracing study^52^. We also identified genes not previously connected to CNCCs that include *Dkk1*, *Foxf1*, *Rgs5*, *Isl2* and *Gata3*. Finally, the third state is SM-CNCCs within the OFT that express *Acta2* smooth muscle genes together with *Gata4*, *Gata5* and *Gata6*. Supporting their requirement, conditional deletion of *Gata6* in NCCs results in septation defects of the OFT^53^. We additionally found novel genes not yet connected to these cells, including *Meox1*, *Bambi*, *Smad6* and *Smad7*. We found that progression of CNCC fate toward smooth muscle of the OFT is associated with progressive downregulation of genes involved in pharyngeal embryonic development and progressive increase in cardiac specification genes, consistent with maturation of the unipotent cellular state to smooth muscle cells (Fig. 8A). Development of NCCs in the pharyngeal apparatus is regulated by cell-cell signaling, in particular from the pharyngeal mesoderm as uncovered by studies of *Tbx1* mutant embryos^34, 54^. In global null or *Mesp1^Cre^* mediated *Tbx1* conditional null embryos, there is altered deployment of CNCCs and reduced contribution to the OFT leading to a persistent truncus arteriosus^34, 54^. Our results suggest that NCCs are produced normally in the neural tube in the absence of *Tbx1,* but they fail to migrate to the caudal pharyngeal arches and OFT and we suggest this is due to disrupted signaling from adjacent mesodermal cells to the NCCs. Here, we identified several receptor-ligand interactions that are disrupted by comparing scRNA-seq data from NCCs and the

**Figure 8:**
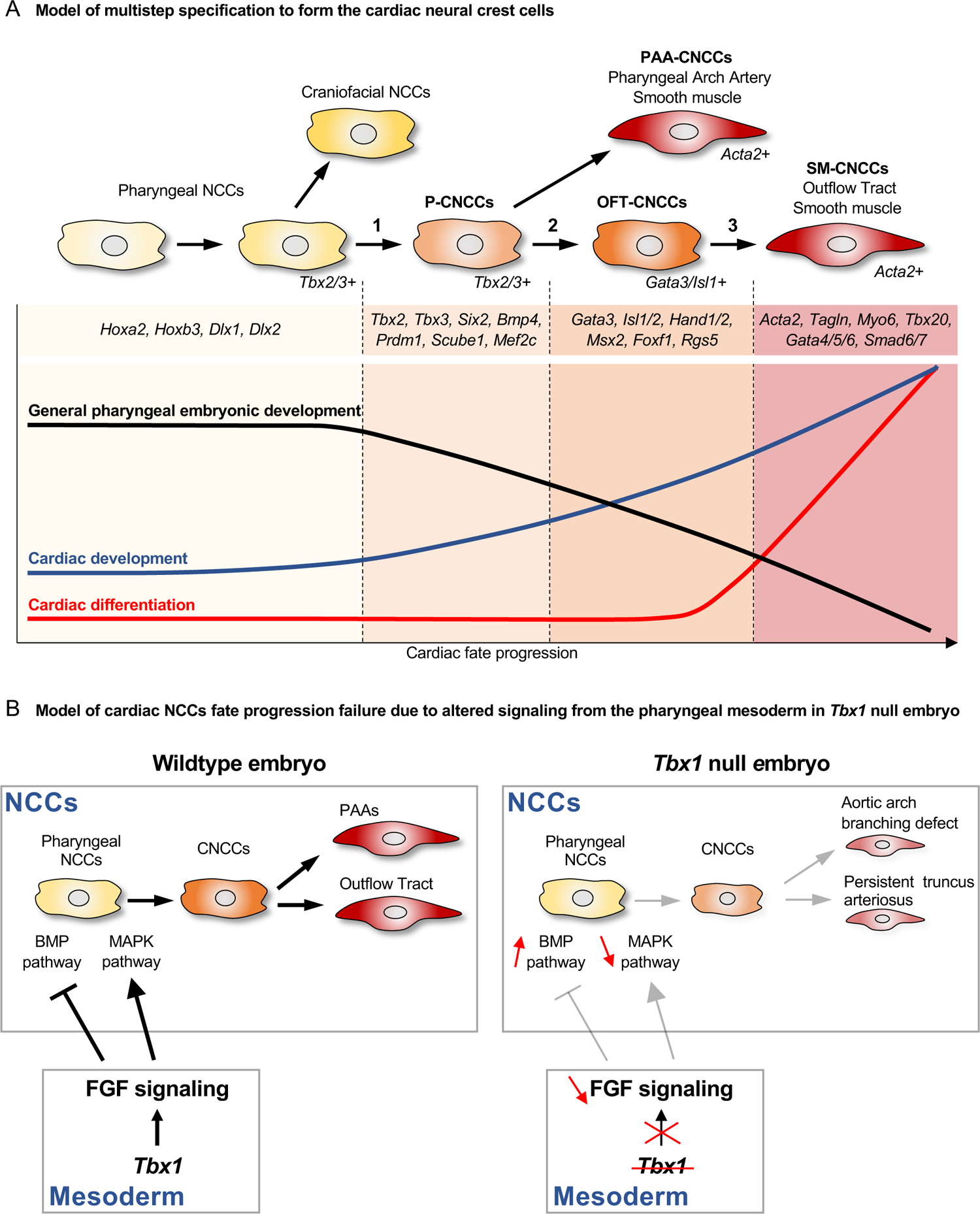
Model of multistep specification to form CNCCs and the signaling pathways disrupted by inactivation of *Tbx1*. A) Colors indicate cell fate progression of pharyngeal NCCs (light yellow) towards vascular smooth muscle cells (red). Step 1 shows the transition between pharyngeal NCCs expressing *Tbx2/3* to cells expressing markers of CNCCs (P-CNCCs). Step 2 shows the transition in which some cells directly differentiate to smooth muscle of PAA-CNCCs, while the majority migrate and enter the OFT as OFT-CNCCs and express *Gata3/Isl1* and Step 3 shows the transition to SM-CNCCs expressing *Acta2*. The graph shows the relative change of biological gene ontology terms during the three transitions of CNCCs to smooth muscle cells. B) Model of non-autonomous signaling between the mesoderm (bottom box) and NCCs (upper box) highlighting the BMP and FGF-MAPK pathways in control (left) and how signaling is disrupted in the absence of *Tbx1* (gray arrows; right). Loss of FGF signaling (red, down arrow) from the mesoderm results in an increase of BMP pathway genes (gray inhibitory arrow; red, up arrow) and decrease of MAPK signaling (gray arrow; red, down arrow) in pharyngeal NCCs. This leads to failure of cardiac fate progression with reduced number of CNCCs and reduced number of SM-CNCCs (gray arrows) with aortic arch/branching defects and failed OFT septation (persistent truncus arteriosus).

### *Mesp1^Cre^* lineages

Using CellChat software^33^, we confirmed known interactions and their disruption in *Tbx1* mutant embryos, now for the first time to NCCs at a single cell level. For example, we identified a reduction of FGF signaling from mesodermal cells to NCCs. Reduced expression of *Fgf* ligands including *Fgf8* and *Fgf10* in the mesoderm of *Tbx1* null embryos have been reported previously^44, 55^. FGF and BMP signaling, both important for NCCs development, can act in an antagonistic manner^48^. In addition, FGF signaling activates the MAPK signaling pathway that is critical for NCC development and migration^31^. Here we propose a model in which FGF paracrine signaling from the mesoderm is required to restrict BMP signaling and activate MAPK signaling in pharyngeal CNCCs necessary for their development and progression to the heart (Fig. 8B). In the absence of *Tbx1*, FGF signaling from the mesoderm is reduced leading to ectopic and overactivation of the BMP pathway and abnormal down-regulation of MAPK pathway in adjacent pharyngeal CNCCs that fail to develop correctly (Fig. 8B). This is consistent with our *in vivo* observation that BMP signaling is abnormally expanded in NCCs in the pharyngeal region of *Tbx1* null mutant embryos and with our *in silico* study showing reduction of downstream effector genes in the MAPK pathway. Reduced phospho-ERK1/2 has previously been reported in NCCs of *Tbx1* null mutant embryos^56^. Our investigation also indicates that BMP4-Bmpr1a+Bmpr2 signaling from mesoderm progenitor populations to NCCs in PA2-6 is abnormally upregulated in *Tbx1* mutant embryos. BMP4 can activate BMP signaling through p-SMAD1/5/9^57, 58^ and our analysis indicates no change in expression of *Bmpr1a* and *Bmpr2* genes in NCCs of *Tbx1* null embryos at E9.5, raising the possibility of a direct upregulation of BMP signaling in CNCCs by mesoderm cells.

Interestingly, our analysis also reveals that Neuregulin (*Nrg1*)-ERBB3 signaling from the MLP to the pharyngeal NCCs of PA2 to PA6 at E9.5 is downregulated in *Tbx1* mutant embryos. Neuregulin is important for migration of NCCs acting as a chemoattractant and chemokinetic molecule^59^ and it is involved in heart development. It has been shown recently that *Nrg1* is a direct transcriptional target gene of *Tbx1* in the multilineage progenitors (MLPs) in the mesoderm^35^. Therefore, alteration of Neuregulin signaling in *Tbx1* mutant embryo could contribute to explain failed cardiac contribution of the NCCs. Together, our analyses indicate that a combination of important signaling from the pharyngeal mesoderm to NCCs are affected in *Tbx1* mutant embryos and could contribute to failure of fate progression of CNCCs.

The pharyngeal endoderm is also an important source of signaling during development, including FGF ligands^60^, that could potentially affect CNCCs development. It will be interesting to evaluate how the exchange of signaling between the pharyngeal endoderm and NCCs are affected in the absence of *Tbx1*.

In conclusion, in this report we identified the transcriptional signature that defines the CNCCs and identified the gene expression dynamics that regulates CNCC fate progression into smooth muscle of the OFT and PAAs. In addition, we highlight direct alteration of FGF signaling from the mesoderm to CNCCs resulting in an abnormal increase in the BMP pathway and failed cardiac contributions in the absence of *Tbx1* at a single cell level. Together our results allow a better understanding of the normal development of CNCCs and provide new insights into the origin of congenital heart defects associated with defective NCCs and 22q11.2DS.

## Methods

### Mouse lines

The following transgenic mouse lines were used: *Wnt1-Cre* ^13^, *ROSA26R-GFP^f/+^* ^14^ that we refer to as *ROSA-EGFP*, *Tbx1^+/-^*^61^, *Tbx1f/+* ^61^. Mice were maintained on a mixed Swiss Webster genetic background. *Tbx2^f/+^* and *Tbx3^f/+^* mutant mouse lines were generated in Dr. Chenleng Cai’s laboratory by inserting a two LoxP sites into the intron sequences flanking exon 2 of the *Tbx2* gene and exons 2-4 of *Tbx3* gene, by gene targeting using homologous recombination. When the genomic sequences between the LoxP sites are floxed out, the reading frame and T-Box domain of *Tbx2* and *Tbx3* are both disrupted. *Tbx2^f/+^* and *Tbx3^f/+^* mice were maintained on a mixed Swiss Webster and C557BL/6 genetic background. Mice and embryos resulting from the different crosses were genotyped by PCR using standard protocols from DNA extracted from toes tips or yolk sac. Animal experiments were carried out in agreement with the National Institutes of Health and the Institute for Animal Studies, Albert Einstein College of Medicine (https://www.einsteinmed.org/administration/animal-studies/) regulations. The IACUC number protocol is #00001034. Embryos were collected at different embryonic days dated from the day of the vaginal plug (E0.5). For each experiment, a representative result is presented from at least three analyzed embryos.

### Immunofluorescence staining on paraffin sections

Embryos were collected in cold 1x PBS (Phosphate Buffered Saline) and fixed in 4% PFA (Paraformaldehyde) for 1 hour at 4°C under constant agitation. Embryos were progressively dehydrated in ethanol then xylene and embedded in paraffin (Paraplast X-tra, Sigma P3808). The embryos were sectioned to 10μm thickness and sections were deparaffinized in xylene and progressively rehydrated in an ethanol series. Sections were incubated and boiled in antigen unmasking solution (Vector laboratories, H-3300) for 15 minutes. After cooling at room temperature, sections were washed in PBS containing 0.05% Tween (PBST) and blocked for 1 hour in TNB buffer (0.1M Tris-HCl, pH 7.5, 0.15M NaCl, 0.5% Blocking reagent [PerkinElmer FP1020]) at room temperature. Then sections were incubated with primary antibodies diluted in TNB overnight at 4°C. Sections were washed in PBST and incubated with secondary antibodies diluted in TNB for 1 hour at room temperature. After washes in PBST, nuclei were stained with DAPI (1/1,000; Thermo Scientific, 62248) and slides were mounted in Fluoromount (Southern Biotech). Embryonic sections were imaged using a Zeiss Axio Imager M2 microscope with ApoTome.2 module. The following primary antibodies were used: goat anti-GFP (1/200, Abcam ab6673), mouse anti-alpha smooth muscle actin ACTA2) (1/200, Abcam ab7817), rabbit anti-αSMA (ACTA2; 1/200, Abcam ab5694), mouse anti-Isl1/2 (1/100, DSHB 40.2D6 and DSHB 39.4D5), mouse anti-TBX2 (1/100, Santa Cruz sc514291), rabbit anti-TAGLN (1/200, Abcam ab14106), rabbit anti-pSMAD1/5/9 (1/100, Cell Signaling D5B10) and rabbit anti-TBX1 (1/100, Lifescience LS-C31179). Donkey secondary antibodies from Invitrogen (Thermo Fisher Scientific) were used (anti-goat, anti-mouse and anti-rabbit).

### RNAscope

#### RNAscope on wholemount embryos

Embryos were collected and dissected in 1x PBS at 4°C and fixed in 4% PFA overnight. Then embryos were dehydrated in progressive methanol washes and stored in 100% methanol at −20°C. Wholemount RNAscope was performed using RNAscope Multiplex Fluorescent Detection Reagents v2 kit (Advanced Cell Diagnostics, ref 323110). Embryos were progressively rehydrated in 1x PBS containing 0.01% Tween (PBST) and were permeabilized using Protease III (Advanced Cell Diagnostics, ref 323110) for 20 minutes at room temperature followed by washes in PBST. Embryos were then incubated with 100μl of pre-warmed mixed C1, C2 and C3 (ratio 50:1:1, respectively) RNAscope probes at 40°C, overnight. After 3 washes in 0.2x SSC (Saline Sodium Citrate), 0.01%Tween, embryos were fixed in 4% PFA for 10 minutes at room temperature. Embryos were then incubated in Amp1 for 30 minutes at 40°C, Amp2 for 30 minutes at 40°C and Amp3 for 15 minutes at 40°C, with washes in 0.2x SSC, 0.01%Tween between each step. Tyramide Signal Amplification (TSA) solutions were prepared as follows: 1/500 for TSA-Fluorescein (Akoya Biosciences, NEL741001KT), 1/2000 for TSA-CY3 (Akoya Biosciences, NEL744001KT) and 1/1000 for TSA-CY5 (Akoya Biosciences, NEL745001KT). To reveal C1 probes, embryos were incubated in HRP1-C1 for 15 minutes at 40°C then washed and incubated with the chosen TSA solution, 30 minutes at 40°C. The amplification reaction was blocked using HRP-Blocker during 15 minutes at 40°C. C2 and C3 probes were revealed following the previous steps and using HRP-C2 for C2 probes and HRP-C3 for C3 probes. Nuclei were stained overnight with DAPI. Wholemount embryos were imaged as Z-stacks using a Leica SP5 confocal microscope or a Nikon CSU-W1 spinning disk confocal microscope. 3D image reconstruction and analyses were performed using Fiji and ImarisViewer 9.8.0 software.

#### RNAscope on cryosections

Embryos were collected and dissected in 1x PBS at 4°C then fixed in PFA 4% overnight and incubated in successive 10%, 20% and 30% sucrose (Sigma-Aldrich S8501) solutions and then embedded in OCT (Optimal Cutting Temperature compound). Embryos were stored at −80°C until they are used. RNAscope was performed on 10μm sections mounted on SuperFrost Plus slides (FisherScientific, 12-550-15) following the RNAscope Multiplex Fluorescent Reagent Kit v2 assay protocol from Advanced Cell Diagnostics.

#### Probes used for RNAscope

*Egfp* (400281, 400281-C3), *Tbx2* (448991), *Hoxb3* (515851), *Dlx2* (555951), *Bambi* (523071), *Hoxa2* (451261), *Gata3* (4033321-C2), *Isl1* (451931-C2), *Tbx3* (832891-C2), *Sox10* (435931-C2), *Msx2* (421851-C2), *Meox1* (530641-C2), *Dlx5* (478151-C3).

### Histology and staining with Hematoxylin & Eosin

Fetuses were collected and dissected in 1x PBS and fixed overnight in 4% PFA. They were progressively dehydrated in ethanol and incubated in xylene prior to embedding in paraffin. Tissue sections of 12μm thickness were deparaffinized in xylene and progressively rehydrated in ethanol washes and incubated for 10 minutes in Hematoxylin (Sigma-Aldrich, HHS16) then rinsed in water and dehydrated in 70% ethanol. Sections were then incubated in alcoholic Eosin (70%) (Sigma-Aldrich, HT110116) solution and progressively dehydrated in ethanol and xylene washes prior to mounting in Permount mounting medium (Fisher Chemical SP15100). Sections were imaged using a Zeiss Axioskop 2 plus microscope.

### Intracardiac India ink injection

Fetuses at E15.5 were dissected in 1x PBS and the chest was carefully opened to avoid damaging the cardiovascular system. A solution containing 50% India ink and 50% 1x PBS was injected into the left ventricle of the heart by blowing gently into an aspirator tube assembly connected to a microcapillary. Immediately after filling the left ventricle and arterial branches, the heart and aortic arch with arterial branches were imaged using a Leica MZ125 stereomicroscope.

### scRNA-seq data generation

Embryos were collected and microdissected in 1x PBS at 4°C. Dissected tissues of interest were maintained in DMEM (Dulbecco’s Modified Eagle Medium, GIBCO 11885084) at 4°C. For E8.5 embryos, the rostral half of the embryos including the heart was collected. At E9.5 and E10.5 the pharyngeal region plus heart were collected. Pharyngeal arch 1 was removed at E10.5 as shown in Fig. 1. Then, tissues were incubated in 1ml of 0.25% Trypsin-EDTA (GIBCO, 25200056) containing 50U/ml of DNase I (Milipore, 260913-10MU), at room temperature for 7 minutes. Next, heat inactivated FBS (Fetal Bovine Serum, ATCC, 30-2021) was added to stop the reaction at a final concentration of 10%, at 4°C. Dissociated cells were centrifugated for 5 minutes at 300 x g at 4°C and the supernatant was removed. Cells were then resuspended in 1x PBS without Ca^2+^ and Mg^2+^ (Coming, 21-031-cv) containing 10% FBS at 4°C and filtered with a 100μm cell strainer. A total of 1μl DAPI (1mM) (Thermo Fisher Scientific, D3571) was added before FACS using a BD FACSAria II system. EGFP positive, DAPI negative cells were then centrifuged at 4°C, 5 minutes at 300 x g and resuspended in 50μl of 1xPBS without Ca^2+^ and Mg^2+^ with 10% FBS. Cells were then loaded in a 10x Chromium instrument (10x Genomics) using the Chromium Single Cell 3’ Library & Reagent kit v2 or single index Chromium Next GEM Single Cell 3’ GEM, Library & Gel Bead kit v3 or v3.1.

### Sequencing

Sequencing of the DNA libraries was performed using an Illumina Hiseq4000 system (Genewiz Company, South Plainfield, NY, USA) with paired-end, 150 bp read length.

### scRNA-seq data analysis

CellRanger (v6.0.1, from 10x Genomics) was used to align scRNA-seq reads to the mouse reference genome (assembly and annotation, mm10-2020-A) to generate gene-by-cell count matrices. All the samples passed quality control measures for Cell Ranger version 6.0.1 (Supplementary Table 1).

#### Seurat analysis for filtering and clustering

Individual scRNA-seq sample data were analyzed using Seurat V4.0.5 ^15^, with parameters as recommended by Seurat software.

#### Integrated scRNA-seq analysis

After individual samples were analyzed by Seurat for clustering, the data were integrated by the RISC software (v1.5) using the Reference Principal Component Integration (RPCI) algorithm for removing batch effects and aligning gene expression values between the control and *Tbx1* null samples at E9.5 and E10.5 ^29^. The integrated data were re-clustered by RISC, using parameters adjusted to match the cell type clusters in the Seurat results. Gene expression differences between control and *Tbx1* null embryos was determined by RISC software for each of the clusters at an adjusted p-value < 0.05 and log2(fold change) > 0.25. The GO enrichment for the differentially expressed genes were identified by clusterProfiler (v4.0.5) ^32^. Cell compositions were computed from the integrated cell clusters and used for two-proportion Z-test as implemented in the R prop.test() function to evaluate the statistical significance in changes between control and null embryos.

#### Cell trajectory analysis

CellRank (v1.5.1) ^21^ was used to infer differentiation trajectory, focusing on determining the probability of cells to adopt the smooth muscle, *Acta2*+ cell fate. The analysis used RNA velocity from Velocyto (v0.17.17) ^62^ and scVelo (v0.2.4) ^62^, and cell-cell similarity to infer trajectories and cell differentiation potential. The analysis was performed for all cells in either the E9.5 control sample or the two E10.5 control samples and then for the cells in the selected clusters that were predicted to have connections to the smooth muscle *Acta2*+ cluster.

#### Cell-cell communication analysis

CellChat (v1.1.3) ^33^ was used to identify the ligand-receptor interactions between *Mesp1^Cre^* mesoderm lineage and NCC lineages and then compare the change between control and *Tbx1* null mutant data at p < 0.05. Data included ligands in the *Mesp1^Cre^* lineage and receptors in CNCC lineage.

## Data Availability

All scRNA-seq datasets generated in this study have been submitted to GEO (Gene Expression Omnibus) repository July 13, 2022, and are awaiting approval.

## Acknowledgements

We thank members of the Genomics core, Flow Cytometry and Analytical Imaging facilities at Albert Einstein College of Medicine. We are grateful to Professor Chenleng Cai for *Tbx2^f/+^* and *Tbx3^f/+^* mice. We thank Professor Robert G. Kelly for insightful comments on the manuscript. This work was supported by grants from the National Institutes of Health (P01HD070454, R01HL138470, R01HL153920, R01HL163667) and by grant from the Fondation Leducq (Transatlantic Network of Excellence 15CVD01). C.D thanks the Fondation Bettencourt-Schueller and the Philippe Foundation for their financial support.

## Author contributions

C.D. and B.E.M. designed the study and experiments. C.D. performed all wet laboratory experiments. C.D., Y.L., A.F., A.V. performed computational analysis of single cell RNA-sequencing data. Y.L., A.F. and D.Z. provided bioinformatics expertise and guidance. C.D. and B.E.M wrote the manuscript. All authors read, intellectually contributed, edited, and approved the manuscript.

---

**Supplementary table 1: Summary of scRNA-seq experiments.** This is summary of the scRNA-seq experiments shown in Figures 1, 2, 3, 5 and 7.

**Supplementary Figure 1:** *Dlx* and *Hox* genes provide proximal-distal and anterior-posterior identities of NCCs in the pharyngeal arches, respectively. A) UMAP plots showing expression levels of *Hoxb3*, *Hoxa2*, *Sox10*, *Dlx2*, *Dlx5* and *Dlx6* genes in cell specific clusters in scRNA-seq data of NCCs at E9.5. B) Wholemount RNAscope *in situ* hybridization of *Wnt1-Cre;ROSA-EGFP* embryos at E9.5 with probes for *Egfp*, *Dlx2* and *Dlx5*. PA, pharyngeal arch. Scale bar: 200 μm. This figure is related to Figure 1.

**Supplementary Figure 2: TBX1 expression is not detected in NCCs at E8.5, E9.5 and E10.5.** Immunostaining for EGFP (green) and TBX1 (red) on sagittal sections of *Wnt1-Cre;ROSA26-EGFP* embryos E8.5 (A) (n=3) and E9.5 (B) (n=6) and on transverse sections of *Wnt1-Cre;ROSA26-EGFP* embryos at E10.5 (C) (n=5). Note that TBX1 is not noticeably expressed in NCCs, but it is expressed in adjacent mesodermal cells. PA, pharyngeal arch; NT, neural tube; end, endoderm; OFT, outflow tract. Scale bars: 100 μm.

**Supplementary Figure 3:** Conditional deletion of *Tbx1* in NCCs does not affect heart development. Hematoxylin and eosin staining on *Wnt1-Cre;Tbx1^f/+^* (n=3) and *Wnt1-Cre;Tbx1^f/f^* (n=3) embryos at E14.5 showing normal aorta and pulmonary trunk septation (A,B) and normal interventricular septation in *Wnt1-Cre;Tbx1^f/f^* embryos (C,D). Ao, aorta; PT, pulmonary trunk; RA, right atrium; LA, left atrium; RV, right ventricle; LV, left ventricle. Scale bars: 500 μm.

**Supplementary Figure 4:** Bubble plots for all ligand-receptor pairs showing significant cell-cell signaling changes from mesodermal cells to NCCs in control and *Tbx1* mutant embryos at E9.5. This version is the complete version compared to the image shown in Fig. 6B.

**Supplementary data 1:** Marker genes and statistics of cell clusters from scRNA-seq of NCCs at E8.5. These data are related to Fig. 1.

**Supplementary data 2:** Marker genes and statistics of cell clusters from scRNA-seq of NCCs at E9.5. These data are related to Fig. 1 and 2.

**Supplementary data 3:** Marker genes and statistics of cell clusters from scRNA-seq of NCCs at E10.5. These data are related to Fig. 1 and 3.

**Supplementary data 4:** Heatmap of gene expression that correlates with cardiac fate probabilities with cells ordered by fate probabilities at E9.5. This heatmap is the complete version of the selected genes shown in Fig. 2.

**Supplementary data 5:** Heatmap of gene expression that correlates with cardiac fate probabilities with cells ordered by fate probabilities at E10.5. This heatmap is the complete version of the selected genes shown in Fig. 3.

**Supplementary data 6: Gene ontology biological processes and disease processes of lineage driver genes at E10.5.** These are the complete lists of gene ontology biological processes and disease processes in each group of genes after dividing the ordered gene list in Supplementary data 5 into four groups of equal number of genes from *Bmp4* to *Gata6*.

**Supplementary data 7:** List of differentially expressed genes and statistics for each NCC cluster of integrated data from control and *Tbx1* null embryos at E9.5. This list is associated with the data shown in Fig. 5B and C.

**Supplementary data 8:** Gene ontology biological processes of upregulated genes in proximal PA2 (C8 in Figure 5) of control and *Tbx1* null embryos at E9.5. This is a complete version of the data shown in Fig. 5F.

**Supplementary data 9:** Gene ontology biological processes of downregulated genes in proximal PA2 (C8 in Figure 5) of control and *Tbx1* null embryos at E9.5. This is a complete version of the data shown in Fig. 5G.

**Supplementary data 10:** List of differentially expressed genes and statistics for each cell cluster of integrated data from control and *Tbx1* null embryos at E10.5. This list is related to data shown in Fig. 7B and C.

**Supplementary data 11:** Gene ontology biological processes of downregulated genes in pharyngeal NCCs (C4 in Figure 7) from control and *Tbx1* null embryos at E10.5. This is a complete version of the data shown in Fig. 7G.

**Supplementary data 12:** Gene ontology biological processes of upregulated genes in OFT-CNCCs (C10 in Figure 7) from control and *Tbx1* null embryos at E10.5. This is a complete version of the data shown in Fig. 7H

