## Supplementary figures and images for "Single cell transcriptomics uncovers a non-autonomous *Tbx1*-dependent genetic program controlling cardiac neural crest cell deployment and progression"

### Supplementary Data

#### Supplementary Figure 1

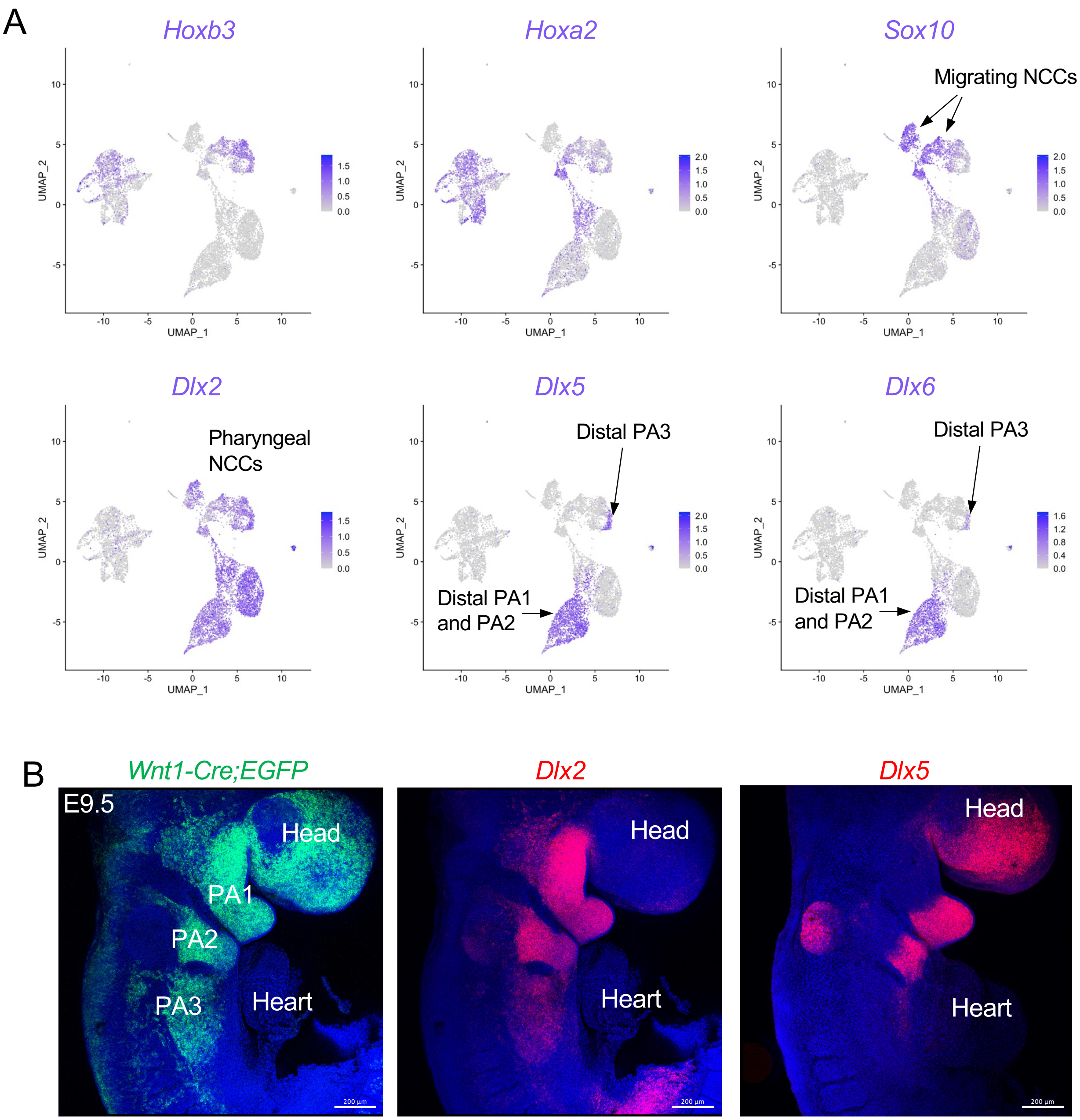

#### Supplementary Figure 2

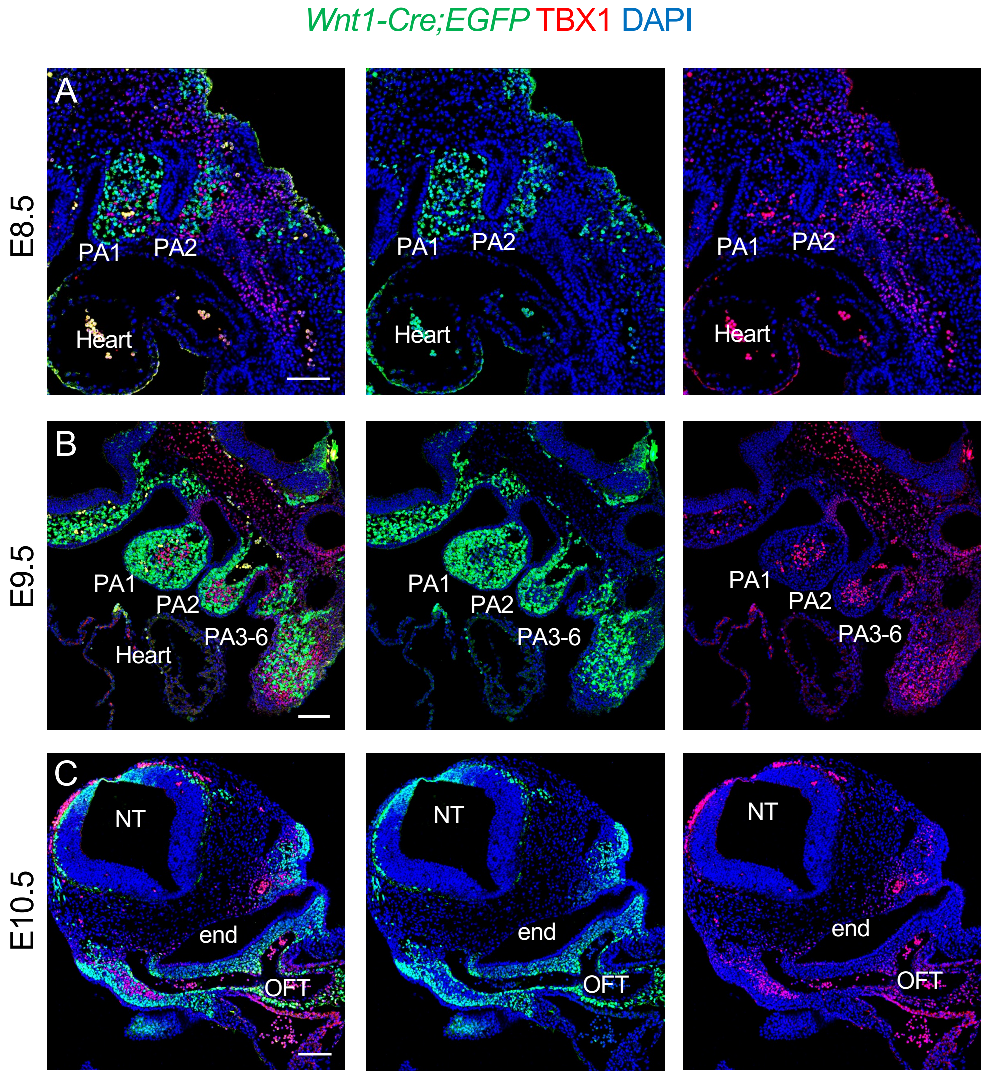

#### Supplementary Figure 3

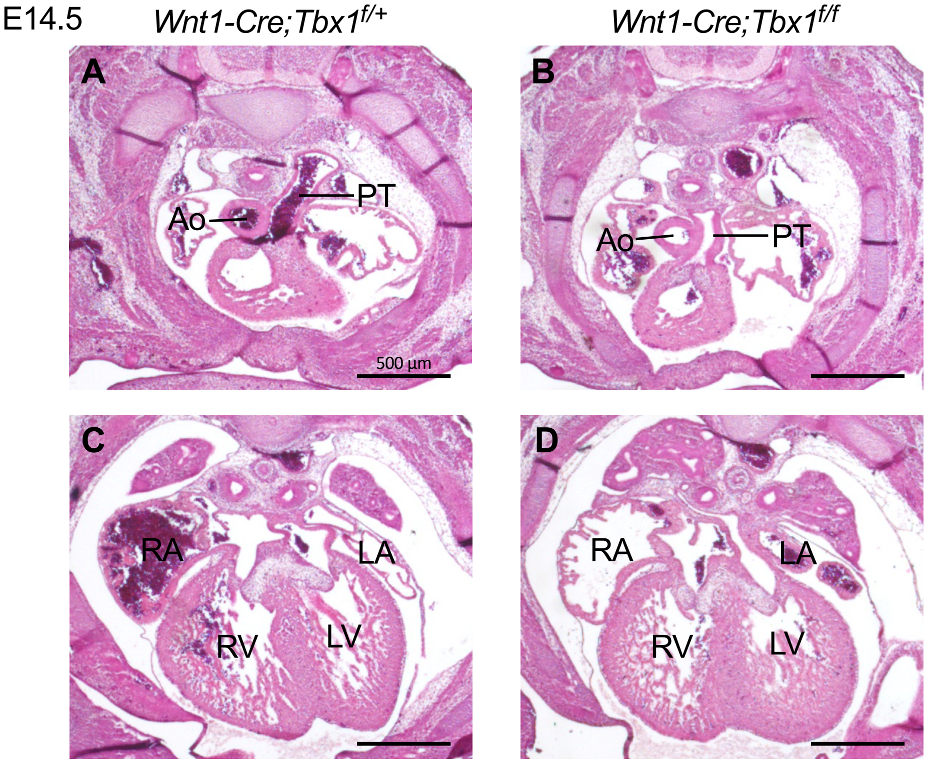

#### Supplementary Figure 4

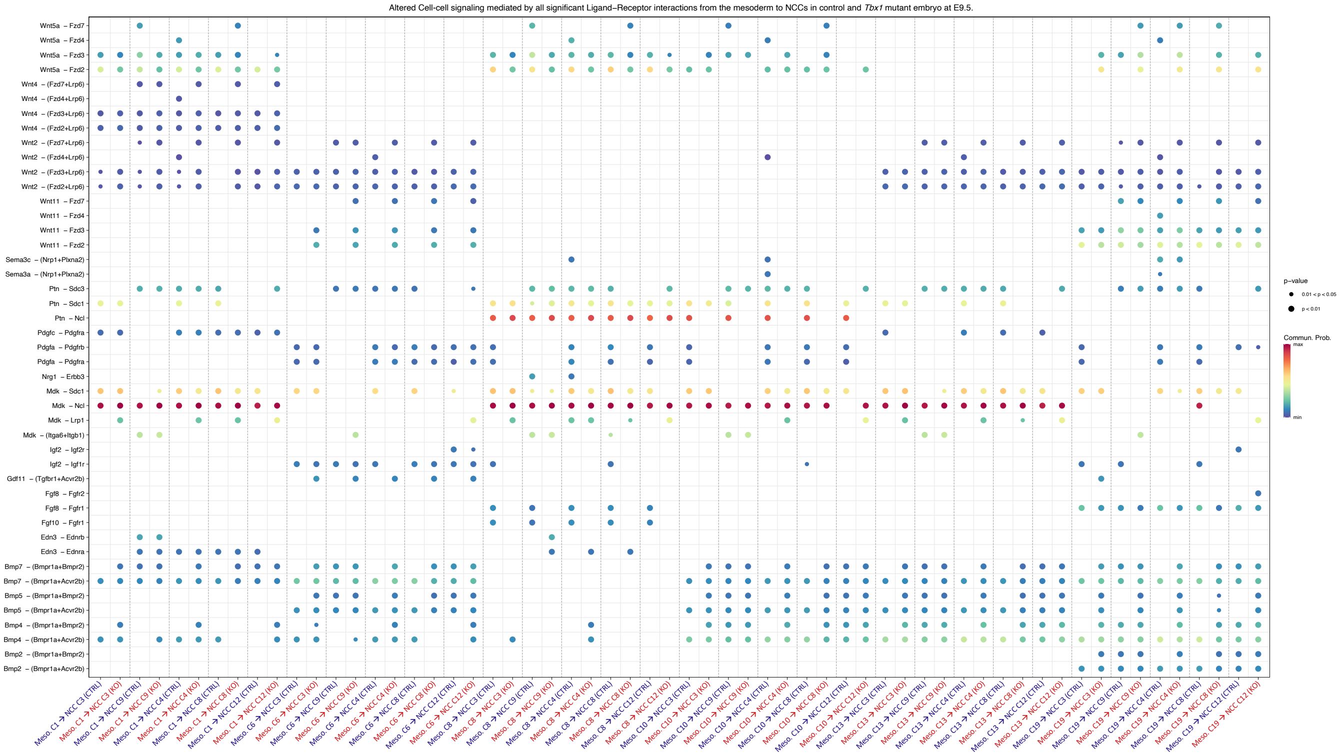
